# Heterogeneous and specific localization of mRNAs at dendritic spine and its dependence on microtubule entry

**DOI:** 10.64898/2026.02.13.705666

**Authors:** Xiaojie Wang, Jiaxing Song, Jiahao Lin, Tian Tian, Alex Tsun-Lok Lui, Bryan Cheuk-Yu Lee, Liang Zhang, Kwok-On Lai

## Abstract

Local mRNA translation near neuronal synapses is crucial for structural plasticity during brain development and memory formation. Whereas mRNAs are transported from soma to dendrites as individual ribonucleoprotein (RNP) granules, how do they subsequently localize at synapses is unclear. In particular, whether different dendritic mRNAs might display differential synaptic localization and interact with specific postsynaptic proteins remains unexplored. Here we found that two well-known dendritic mRNAs, *Actb* and *Camk2a,* are targeted to different subdomains of dendritic spines and delivered to different spine populations after synaptic stimulation. Surprisingly, the induced mRNA localization to dendritic spines is associated with microtubule (MT) entry and specifically requires the MT-associated motor KIF5A. Proximity-labelling proteomics further identifies distinct postsynaptic proteins enriched by *Actb* and *Camk2a*, from which we uncover the cytoskeletal protein Septin7 to be co-localized with the two mRNAs at different subdomains of dendritic spines. Our study reveals an unexpectedly specific mRNA localization mechanism at synapses and demonstrates proximity-labelling as a useful approach to map their synaptic partners.

## INTRODUCTION

Neurons extend dendritic arbors that span hundreds of micrometers from the soma, creating a spatial challenge for the distal synapses to maintain their highly specific molecular compositions. To help tackling this problem, neurons develop a two-way communication between synapses and the nucleus: first, synaptic activity regulates transcription via calcium signaling and synapse-to-nucleus transport of signaling molecules (Greer and Greenberg, 2008; Lai et al., 2008; Andres-Alonso et al., 2023). Second, some of the transcripts are transported to the synapses for local translation, which enables rapid changes of their structure and function in response to synaptic stimulation (Doyle and Kiebler, 2011). This local mRNA translation is fundamental to consolidating long-term synaptic plasticity and memory (Steward and Schuman, 2001; Holt et al., 2019; Cajigas et al., 2012; Buxbaum et al. 2014). The long-range soma-to-dendrite transport of mRNAs requires the recruitment of specific RNA-binding proteins (RBPs) to form the RNP granules, which are then carried along the microtubules by kinesin motor proteins (Kanai et al., 2004; Kiebler and Bassell, 2006; Mitsumori et al., 2017). To achieve synapse-specific plasticity, mRNA translation can occur within a single dendritic spine, thereby selectively modulating the protein composition of the activated postsynaptic sites (Doyle and Kiebler, 2011). Chemical long-term potentiation (cLTP) increases the abundance of mRNAs such as *Camk2a*, *Actb*, and *Psd95* near dendritic spines (Donlin-Asp et al., 2021). Stimulation of individual spines using two-photon glutamate uncaging also recruits *Rgs4* and *Actb* mRNA granules to the activated spines (Yoon et al., 2016; Bauer et al., 2019), indicating the presence of an activity-dependent mechanism to capture mRNAs at the synapses.

Compared to the microtubule-mediated long-range transport, much less is known about the short-range delivery and subsequent localization of mRNAs to the dendritic spines. The actin/myosin motor system has long been regarded as the major short-range transport system in neurons (Hammer III and Sellers, 2011). However, synaptic stimulation can drive the entry of microtubule and its associated proteins to dendritic spines (Jaworski et al., 2009; Hu et al., 2011), raising the possibility that kinesin might also contribute to the short-range synaptic delivery of molecules. Instead of being transported together as single entities, individual mRNAs are carried to the dendrites as separate RNP granules (Tübing et al., 2010; Mikl et al., 2011; Batish et al., 2012) which contain diverse components of both mRNAs and proteins (Fritzsche et al., 2013; Mitsumori et al., 2017). Recent multiplexed *in situ* hybridization also reveals that thousands of different dendritic mRNAs are heterogeneously distributed along the dendrite (Wang et al., 2020). However, the key question of whether these initially separate RNP transport granules end up at the same or different dendritic spines as destinations after synaptic stimulation remains unanswered. Given that diverse protein compositions can be observed among individual synapses of the same neuron (Grant and Fransen, 2020; Oostrum and Schuman, 2024) and different proteins can be distributed at specific subdomains within a single spine, it is important to investigate whether mRNAs undergo similar diversity in their synaptic targeting. Furthermore, how is the specificity in synaptic mRNA localization achieved? Do mRNAs localize at dendritic spines through interaction with specific postsynaptic proteins?

To begin addressing these questions, we focus on *Actb* and *Camk2a* — two abundant dendritic mRNAs whose encoded proteins have well-characterized synaptic functions in neurons. Through cLTP induction followed by single-molecule fluorescence *in situ* hybridization (smFISH), we found that their delivery to dendritic spines is not random but highly specific. We further uncover the essential role of microtubule entry as well as the kinesin motor KIF5A in the synaptic targeting of mRNAs. The specificity of *Actb* and *Camk2a* localization at dendritic spine is further supported by their enrichment of unique postsynaptic proteins in proximity labelling proteomics. Our findings indicate the existence of a specific synapse-localization mechanism for mRNAs, which represents a previously unidentified post-transcriptional regulation that can fine-tune the composition of individual dendritic spines in response to synaptic stimulation.

## RESULTS

### Differential localization of *Actb* and *Camk2a* mRNAs at specific subdomains of dendritic spines

Both the base and the head of dendritic spine have been reported to contain mRNAs (Tiruchinapalli et al., 2003; Kao et al., 2010; Dynes and Steward, 2012). To ask whether two different mRNAs show preference to be localized at different subdomains of dendritic spine, we perform smFISH with probes coupled to different fluorescence dyes to simultaneously visualize *Actb* and *Camk2a* mRNAs and directly compare their distribution within the same hippocampal neurons. A sense probe was used as negative control to rule out any non-specific signal. Specificity of the smFISH probes were validated through exogenous expression of the transcripts in HEK 293T cells (Fig. S1). We classify the mRNA distribution into three different dendritic domains: (i) the dendritic shaft, (ii) the spine base which represents the area directly under a dendritic spine (Chu et al., 2019; Bimbi and Tongiorgi, 2024), and (iii) the spine head (Fig. 1A). Both mRNAs were more frequently detected at the base than the head of dendritic spines, but a higher percentage of *Actb* was detected in the spine heads compared to *Camk2a* within the same dendrites (Fig. 1B). Consistent with previous studies (Mikl et al., 2011; Batish et al., 2012), there was little co-localization between the two mRNAs on the dendrites. Interestingly, their co-localization was significantly higher at the base of dendritic spines (Fig. 1C-D). In contrast, the two mRNAs rarely colocalized within the same dendritic spine head. This finding suggests that although the two mRNAs are initially transported to the dendrites as separate RNP granules, they are commonly destined to the base of dendritic spine which serves as the hot spot for mRNA localization. To verify this notion, we examined *Actb* mRNA movement using the MS2 system followed by time-lapse imaging (one frame per 10 seconds for 30 minutes). Neurons were co-transfected with the MS2-*Actb* mRNA reporter, GFP-MCP, and tdTomato to label dendritic spines. Compared to the granules on the dendritic shaft, *Actb* mRNAs showed reduced movement at the base of dendritic spines (Fig. 1E-F). Notably, the *Actb* mRNA granules were observed to selectively stop underneath dendritic spines (Fig. 1G). Quantification revealed that more than 40% of the moving *Actb* mRNAs demonstrated selective docking at the spine base (Fig. 1H). These findings confirm the base of dendritic spines plays a crucial role for mRNA localization.

**Figure 1.**
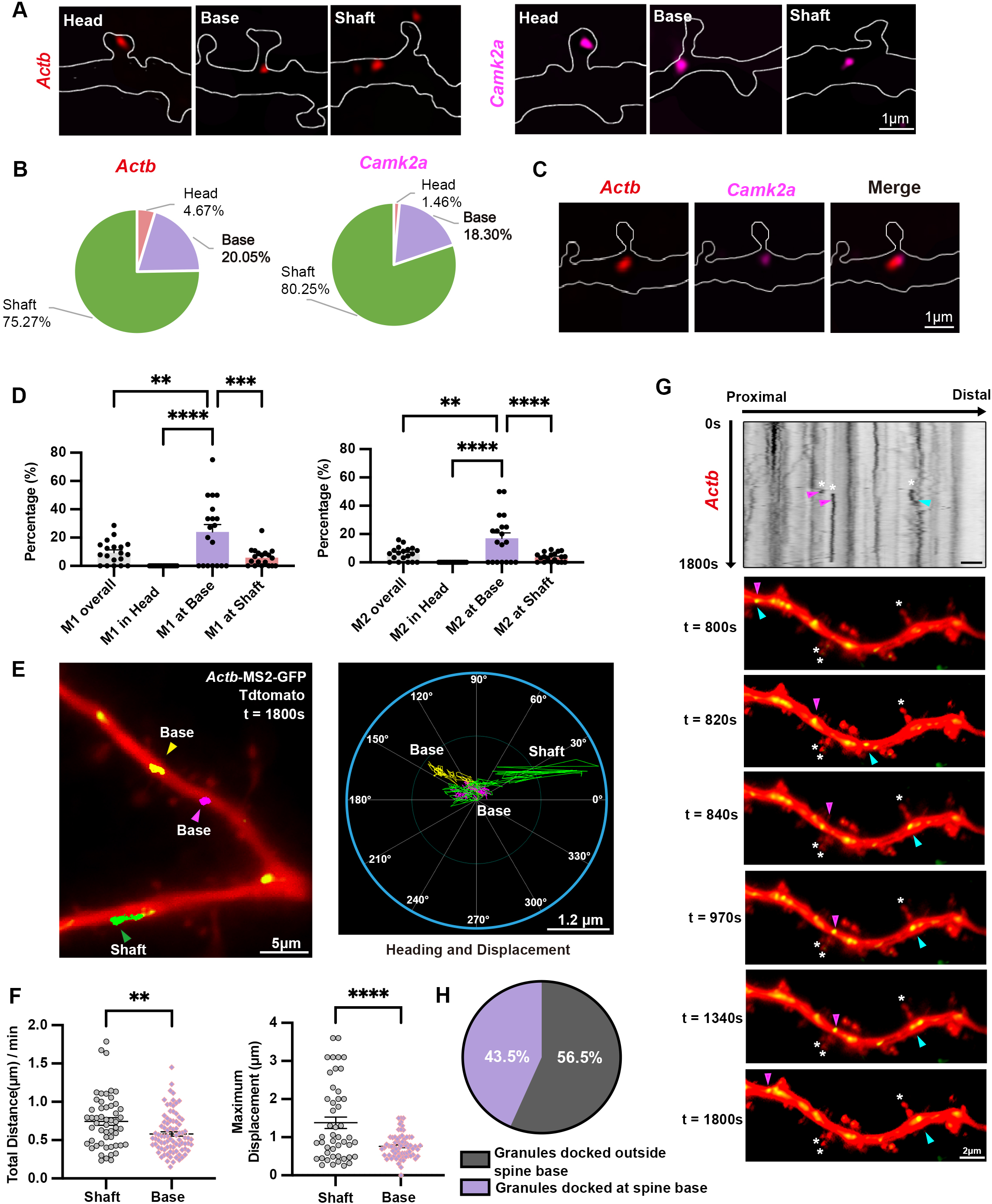
Spine base serves as a hotspot for mRNA localization. (A) Representative dual-color smFISH images of *Actb* (red) and *Camk2a* (magenta) mRNA puncta in different region of dendrites in GFP-transfected rat hippocampal neurons (21 DIV). Dendrites and dendritic spines were outlined based on GFP signals. (B) Both *Actb* and *Camk2a* are more frequently located at the spine base than the spine head. Pie charts showing the percentages of *Actb* and *Camk2a* mRNA puncta distributed to the different dendritic compartments (dendritic shaft, spine base, and spine head). Results were pooled from two independent experiments; 19 neurons were imaged and 364-481 mRNA puncta from 40 primary basal dendrites were analyzed. (C) Representative dual-color smFISH images showing co-localization of *Actb* (red) and *Camk2a* (magenta) mRNAs at the spine base. (D) *Actb* and *Camk2a* mRNAs exhibit preferential co-localization at the spine base. mRNA co-localization was represented by M1 (percentage of *Actb* puncta overlapping with *Camk2a*) and M2 (percentage of *Camk2a* puncta overlapping with *Actb*). Each data point represents one neuron. Results were pooled from two independent experiments; 19 neurons were imaged and 364-481 mRNA puncta from 40 primary basal dendrites were quantified. Data are mean ± SEM; *\*\*p < 0.01, ***p < 0.001, ****p < 0.0001*; one-way ANOVA with Tukey’s multiple comparisons test. (E) Live imaging of MS2-*Actb* mRNA reporter (green) in dendrites of hippocampal neurons co-expressing tdTomato (red). Neurons were transfected at 13 DIV and imaged at 19-21 DIV (one frame/10 s for 1,800 s). Left: trajectories of *Actb* mRNA puncta localized at the spine base (yellow and purple arrowheads) or on the dendritic shaft (green arrowhead) were shown over time. Right: polar plots of heading and displacement for each mRNA. (F) The mRNAs at the spine base show less movement compared to that on the dendritic shaft. Quantification of total distance of movement (left) and maximum displacement (right) for *Actb* mRNAs initially located in the shaft or at the spine base. Results were pooled from two independent experiments; 31 neurons were imaged and 53-84 puncta from 35 dendrites were quantified. Data are mean ± SEM; *\*\*p < 0.01, ****p < 0.0001*; Student’s t-test. (G) Representative images and kymograph showing the docking of fast-moving *Actb* mRNAs (purple and blue arrows) at the base of dendritic spines (white asterisks). Time was indicated in seconds (s). (H) Analysis of fast-moving *Actb* granules (velocity > 0.13 μm/s) to determine where they stop. Near half of the granules selectively docked at the spine base. Results were pooled from two independent experiments; 31 neurons were imaged and 10-13 docking granules from 35 dendrites were quantified.

Next, we investigate the possibility of differential localization of *Actb* and *Camk2a* mRNAs after synaptic stimulation. cLTP is a widely used experimental paradigm to induce activity-dependent plasticity at glutamatergic synapses of cultured hippocampal neurons. The NMDA receptor co-agonist glycine was applied to rat hippocampal neurons to induce cLTP. Thirty minutes after the brief glycine stimulation, there was a significant increase in dendritic spine size, particularly the mushroom spines (representing the mature spines) (Fig. 2A-B, Fig. S2B), which was accompanied by a reduction in the density of the immature filopodia (Fig. S2A). The change in spine morphology therefore validates the effectiveness of the stimulation paradigm. After cLTP, the co-localization of *Actb* and *Camk2a* at the spine base was significantly decreased (Fig. 2C), suggesting that the two different mRNA granules become separate and targeted to different destinations. Notably, the differential distribution of *Actb* and *Camk2a* at the subdomains of dendritic spines becomes more prominent after cLTP, with *Actb* showing increasing localization at the spine heads over time, while *Camk2a* concurrently accumulating at the spine base (Fig. 2D). Our findings therefore reveal that although there is increased localization of both *Actb* and *Camk2a* mRNAs towards dendritic spines after synaptic stimulation, the two different mRNAs display heterogeneous localization at different subdomains of dendritic spines within the same neurons.

**Figure 2.**
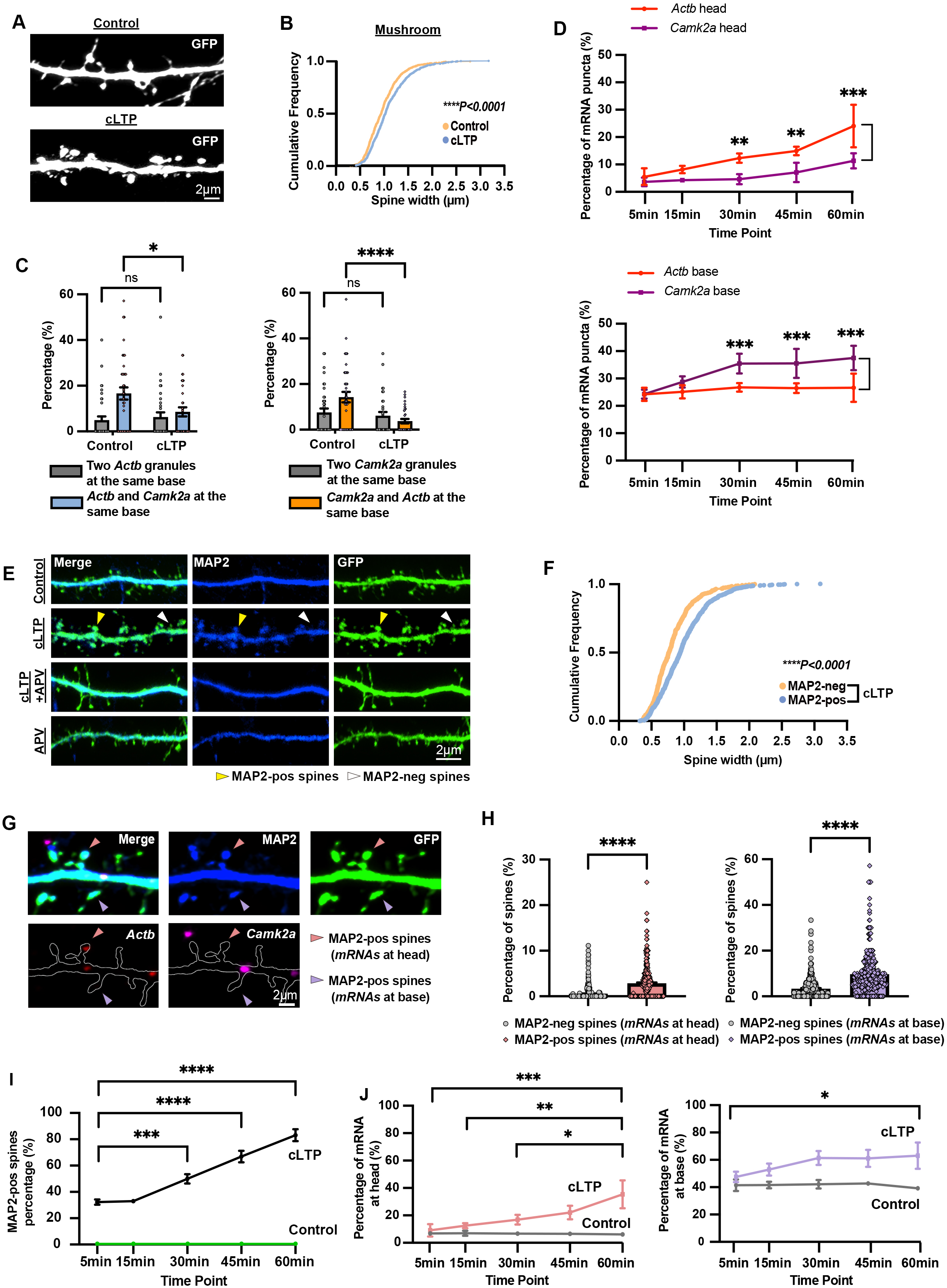
Differential targeting of *Actb* and *Camk2a* mRNAs to dendritic spines in response to cLTP and its correlation with MAP2 entry. (A) cLTP induces spine enlargement. Representative dendritic segments of rat hippocampal neurons (21 DIV) transfected with GFP (13 DIV) and stimulated by cLTP (200 μM glycine and 1 μM strychnine, 6 min) or control treatment. After recovery in conditioned medium for 30 min, neurons were stained with anti-GFP antibody. (B) Cumulative frequency curve shows the width of the mushroom spines under control and cLTP conditions. The spine width significantly increased after the synaptic stimulation. Results were pooled from three independent experiments; 1,757-2,045 spines from 32–37 neurons per condition were analyzed, from which the width of mushroom spines was quantified. Data are mean ± SEM; ****p < 0.0001; Student’s t-test. (C) Co-localized mRNAs at the spine base are relocated after cLTP. Co-presence between the same-(*Camk2a*-*Camk2a* or *Actb*-*Actb*) or different- (*Camk2a*-*Actb*) mRNA granules at the spine base under control and cLTP conditions was quantified. The co-presence of *Actb* and *Camk2a* at the spine base was significantly reduced 30 mins after cLTP. Each data point represents one neuron. Results were pooled from three independent experiments; 32-37 neurons per condition were imaged and 816-1,122 mRNA puncta from 96-111 dendrites per condition were quantified. Data are mean ± SEM; *p < 0.05, ****p < 0.0001; two-way ANOVA with Šídák’s multiple comparisons test. (D) Differential distribution of *Actb* and *Camk2a* mRNA granules at dendritic spine after synaptic stimulation. The proportion of *Camk2a* and *Actb* mRNAs localized at the base and head of dendritic spines across different time points following cLTP was measured. Hippocampal neurons (transfected with GFP at 13 DIV) were subjected to cLTP at 21 DIV, fixed at the indicated time points, and processed for dual color smFISH combined with anti-GFP and anti-MAP2 immunostaining. The localization of mRNA puncta within primary basal dendrites was quantified and categorized as in the shaft, base, or head of dendritic spines. After stimulation, the proportion of *Actb* mRNA in the spine heads increased significantly over time compared to *Camk2a*, whereas the proportion of *Camk2a* mRNA at the spine base showed a significant increase compared to *Actb*. Results were pooled from three independent experiments; 24 neurons were imaged per condition and 541–777 mRNA puncta from 72 primary basal dendrites were quantified per condition. Data are mean ± SEM; **p < 0.01, ***p < 0.001; two-way ANOVA with Šídák’s multiple comparisons test. (E) MAP2 entry to dendritic spine heads after cLTP depends on NMDA receptors. Representative dendritic segments of rat hippocampal neurons from four experimental groups: control, cLTP, cLTP+APV, and APV. Neurons were transfected with GFP at 13 DIV and subjected to the respective treatments at 21 DIV. After recovery for 60 min, neurons were stained with anti-GFP and anti-MAP2 antibodies. Yellow arrows indicate MAP2-positive spines; white arrows indicate MAP2-negative spines. (F) MAP2-positive spines have increased spine width compared to MAP2-negative spines. Cumulative frequency distribution of the width of MAP2-positive (MAP2-pos) and -negative (MAP2-neg) spines. Results were pooled from three independent experiments; 1,200-1,757 spines from 32 neurons per condition were quantified. Data are mean ± SEM; *\*\*\*\*p < 0.0001*; Student’s t-test. (G) Representative dendrites of hippocampal neurons indicating the localization of *Actb* and *Camk2a* mRNAs at MAP2-positive spines after cLTP. Neurons (21 DIV) transfected with GFP (13 DIV) were subjected to cLTP recovered for 30 min before dual-color smFISH and immunostaining with GFP and MAP2 antibodies. MAP2-positive spines with mRNA accumulation at the head (pink arrowhead) or base (purple arrowhead) were indicated. (H) After cLTP, mRNAs preferentially accumulate at the head and base of MAP2-positive spines. Percentage of dendritic spines with either *Actb* or *Camk2a* mRNA localized at the head or base of MAP2-positive and MAP2-negative spines was quantified. Results were pooled from three independent experiments; 32 neurons were imaged, with 2,208–2,441 mRNA puncta and 6,007–7,032 spines quantified. Data are mean ± SEM; *\*\*\*\*p < 0.0001*; Student’s t-test. (I) The proportion of MAP2-positive spines increases over time following cLTP induction. Rat hippocampal neurons were transfected with GFP (13 DIV) and subjected to cLTP stimulation at 21 DIV. Neurons were returned to conditioned medium for the indicated durations (5, 15, 30, 45, 60 min) before fixation. Results were pooled from three independent experiments; 951–1,837 spines from 24-26 neurons per condition were quantified. Data are mean ± SEM; *\*\*\*p < 0.001, ****p < 0.0001*; one-way ANOVA with Tukey’s multiple comparisons test. (J) mRNA localization at the base and head of dendritic spines increases over time following cLTP. Rat hippocampal neurons were transfected with GFP (13 DIV) and subjected to cLTP stimulation at 21 DIV. Neurons were returned to conditioned medium for the indicated durations (5, 15, 30, 45, 60 min) before fixation for dual-color smFISH combined with anti-GFP and anti-MAP2 immunostaining. The number of *Actb* and *Camk2a* mRNA puncta within primary basal dendrites was quantified and categorized as in the dendritic shaft, base, or head of dendritic spines. Results combining both mRNAs were pooled from three independent experiments; 24-26 neurons per condition were imaged, and 1,455-1,914 mRNA puncta from 72 primary basal dendrites were quantified. Data are mean ± SEM; *\*p < 0.05, **p < 0.01, ***p < 0.001*; one-way ANOVA with Tukey’s multiple comparisons test.

### *Actb* and *Camk2a* mRNAs are targeted to different populations of dendritic spines after cLTP

To achieve local translation, mRNAs are packaged into granules and transported along microtubules within the dendrite (Das et al., 2021). MAP2, a microtubule-stabilizing protein, participates in activity-dependent synaptic plasticity, and cLTP results in the invasion of MAP2 from the dendritic shaft to the heads of dendritic spines (Kim et al., 2020). Consistent with this study, we found that stimulation of neurons by cLTP resulted in the presence of MAP2 in dendritic spines after 30 mins (Fig. 2E). Increasing the duration of glycine stimulation (from 3 to 6 minutes) significantly enhanced the proportion of neurons that exhibit MAP2 signals in the spine heads (Fig. S2C). The MAP2 entry to spine was completely abolished by the NMDA receptor antagonist APV, indicating the involvement of NMDA receptors (Fig. 2E, Fig. S2D). For those neurons that showed MAP2 immunoreactivity in the spine heads after cLTP, only about half of their spines demonstrated MAP2 invasion (Fig. S2E), and these MAP2-positive spines were significantly larger than the other spines that were devoid of MAP2 fluorescence (MAP2-negative spines) (Fig. 2F). Notably, the MAP2-positive spines were more likely to capture *Actb* and *Camk2a* mRNAs at their heads and base as compared to the MAP2-negative spines of the same neurons (Fig. 2G-H). Furthermore, after cLTP induction there was a parallel increase between the entry of MAP2 to the spine heads and localization of the two mRNAs at dendritic spines over time (Fig. 2I-J). cLTP did not change the density of either *Actb* or *Camk2a* granules on the dendrites (Fig. S2F). Next, we investigate whether *Actb* and *Camk2a* mRNAs show differential targeting to the two populations (MAP2-positive versus MAP2-negative) of dendritic spines. Towards this end, we focus on the mRNAs within the spine head. Intriguingly, *Actb* and *Camk2a* display differential preferences towards the two populations of spines: *Camk2a* mRNAs in the spine heads were almost exclusively (99%) found in the MAP2-positive spines; in contrast, considerable proportion (∼36%) of *Actb* mRNAs were localized to the heads of MAP2-negative spines (Fig. 3A-B).

**Figure 3.**
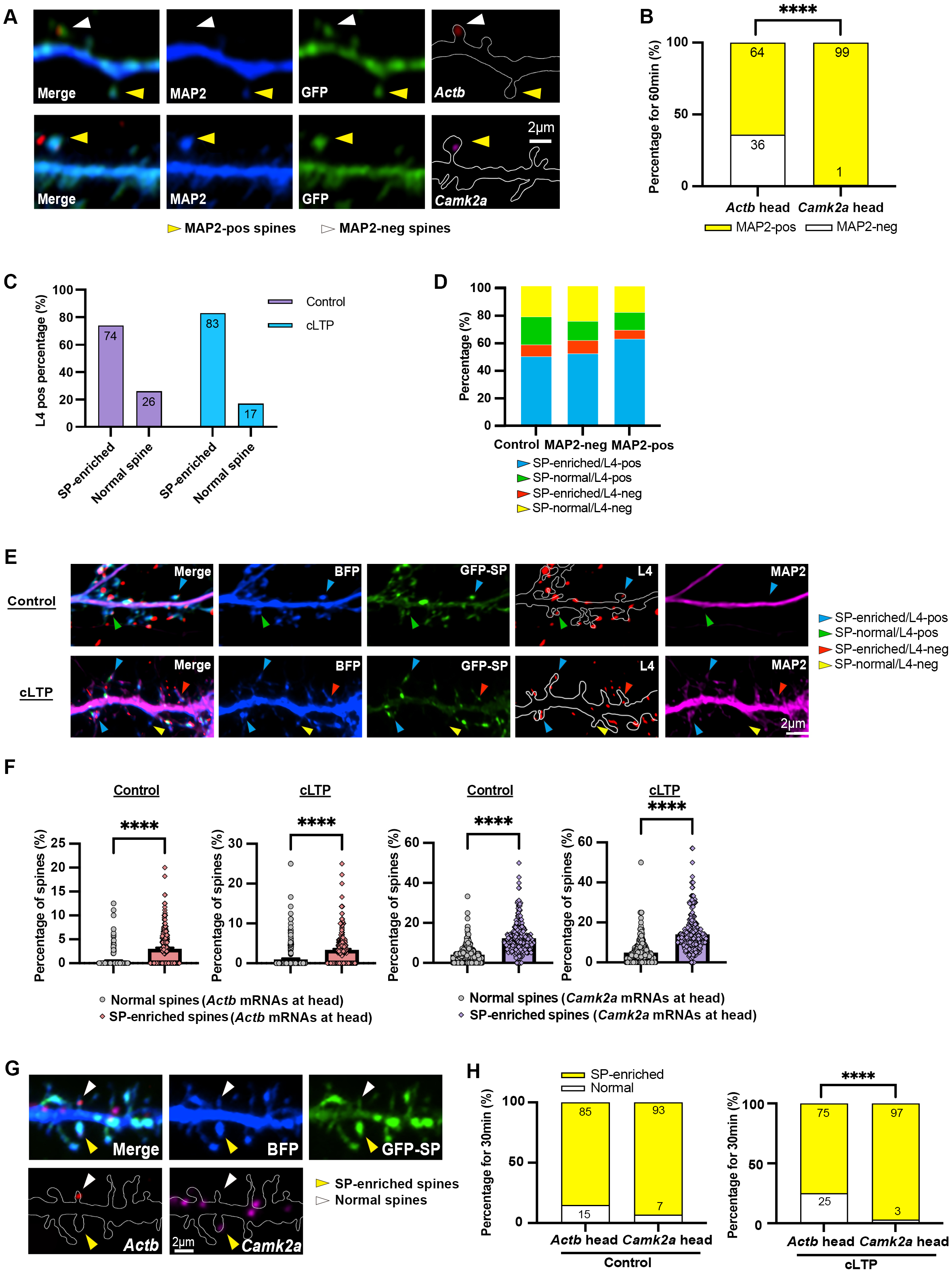
*Actb* and *Camk2a* mRNAs show specific targeting to different populations of dendritic spine heads after cLTP. (A) Representative dendritic segments of rat hippocampal neurons showing the localization of *Camk2a* and *Actb* mRNAs in the heads of MAP2-positive (yellow arrowhead) and MAP2-negative spines (white arrowhead), respectively. Hippocampal neurons transfected with GFP (13 DIV) were subjected to cLTP for 6 mins (21 DIV). After recovery for 60 min, neurons were processed for dual color smFISH combined with immunostaining for GFP and MAP2. (B) *Camk2a* exhibits much stronger preference to localize at the head of MAP2-positive spines than *Actb* after cLTP. Results were pooled from three independent experiments; 32 neurons per condition were imaged and 2,208-2,441 mRNA puncta, 6,007-7,032 spines per condition were quantified. ****p < 0.0001; chi-square test. (C) Higher proportion of synaptopodin (SP)-enriched spines contain ribosomal protein L4. Results were combined from three independent experiments; 3,197–3,598 spines from 28–30 neurons per condition (control or cLTP) were quantified. (D) Upon cLTP induction, higher proportion of MAP2-positive spines are enriched with SP and contain ribosomal protein L4 as compared to the MAP2-negative spines from the same neurons. The latter show similar proportion to control neurons without cLTP. Results were pooled from three independent experiments; 894-1,997 spines from 27-29 neurons were quantified for each experimental condition. (E) Representative images of primary basal dendrites of hippocampal neurons (21 DIV) showing the localization of GFP-SP and ribosomal protein L4 in MAP2-psotive or MAP2-negative spines. Neurons were transfected with BFP and GFP-SP (13 DIV) and subjected to cLTP (or control treatment) at 21 DIV. After recovery for 30 min, neurons were stained with anti-BFP, anti-L4, and anti-MAP2 antibodies. Blue arrowhead indicates SP-enriched, L4-positive spine; red arrowhead indicates SP-enriched, L4-negative spine; green arrowhead indicates SP-normal, L4-positive spine; yellow arrowhead indicates SP-normal, L4-negative spine. (F) *Actb* and *Camk2a* mRNAs prefer to localize at SP-enriched spines. Results were pooled from three independent experiments; 28-30 neurons were imaged for each condition and 1662-1974 mRNA puncta, 3197-3598 spines were quantified. Data are presented as mean ± SEM, *\*\*\*\*p < 0.0001*, Student’s *t*-test. (G) Representative dendritic segments of rat hippocampal neurons showing the localization of *Camk2a* and *Actb* mRNAs in the heads of SP-enriched (yellow arrowhead) and non-SP-enriched (normal spine, white arrowhead) spines, respectively. Hippocampal neurons transfected with GFP (13 DIV) were subjected to cLTP for 6 mins (21 DIV) and processed for dual color smFISH with anti-BFP immunostaining. (H) *Camk2a* exhibits stronger preference to localize at the head of SP-enriched spines. Results from combined from three independent experiments; 28–30 neurons per condition were imaged and 1,662–1,974 mRNA puncta; 3,197–3,598 spines per condition were quantified. *\*\*\*\*p < 0.0001*; chi-square test.

Dendritic spines may also be classified based on their enrichment of specific organelles. For example, stacks of smooth endoplasmic reticulum form a distinctive organelle called the spine apparatus, which is present in subsets of dendritic spines (Konietzny et al., 2023). The spine apparatus is typically located near the spine neck and is believed to regulate calcium homeostasis and protein synthesis within dendritic spines (Pierce et al., 2000, 2001; Segal et al., 2010; Konietzny et al., 2023; Falahati et al., 2025). The actin-binding protein Synaptopodin (SP) is commonly used as a marker for the spine apparatus (Vlachos, 2012). By transfecting hippocampal neurons with BFP and GFP-SP (Fig. S3A), we found that almost half of the dendritic spines were enriched with GFP-SP (SP-enriched spines, 51.75% ± 0.85%, n = 30), suggesting the presence of spine apparatus in these spines. The SP-enriched spines were more likely to contain the ribosomal protein L4 (Fig. 3C), particularly in the MAP2-positive spines after cLTP (Fig. 3D-E). smFISH further revealed that, similar to the MAP2-positive spines, the SP-enriched spines were more likely to contain mRNA granules at their base and head than those without SP enrichment (Fig. 3F, Fig. S3B). These findings prompt us to explore whether *Actb* and *Camk2a* mRNAs also exhibit differential targeting to the SP-enriched spines. Indeed, compared to *Actb*, *Camk2a* displayed a higher preference to the SP-enriched spines, a trend that became more pronounced after cLTP (Fig. 3G-H). Together with the differential preference to MAP2-positive spines (Fig. 3A-B) and presence at different subdomains of dendritic spines (Fig. 2D), our findings strongly suggest that the targeting of *Actb* and *Camk2a* mRNAs to dendritic spines after synaptic stimulation is heterogeneous and highly specific.

### Microtubules are involved in delivering mRNAs to dendritic spines upon synaptic stimulation

Next, we investigate the mechanism by which cLTP induces *Actb* and *Camk2a* mRNAs to dendritic spines. Since dendritic spines that contain MAP2 after synaptic stimulation are more likely to capture mRNAs (Fig. 2H), we reason that microtubule entry plays a crucial role in delivering the two mRNAs to the dendritic spines. To test this hypothesis, we first examine microtubule invasion to dendritic spine by performing a time-course analysis following cLTP on the localization of MAP2 and the microtubule plus-end binding protein EB3. We observed a distinct temporal pattern, in which EB3 presence in the spine head peaked as early as 5 minutes post-stimulation and subsequently declined, whereas MAP2 accumulation in spines increased progressively over time (Fig. 4A-B). This dynamic is consistent with their respective functions: EB3 marks the actively polymerizing microtubule ends that spearheads the initial invasion, while MAP2 entry subsequently stabilizes the newly formed microtubule tracks. Importantly, EB3 granules were specifically enriched in MAP2-positive spines following cLTP (Fig. 4C), demonstrating a strong correlation between dynamic microtubule entry and the formation of a stable, MAP2-rich spine compartment that would be crucial for the subsequent recruitment of mRNAs to these spines.

**Figure 4.**
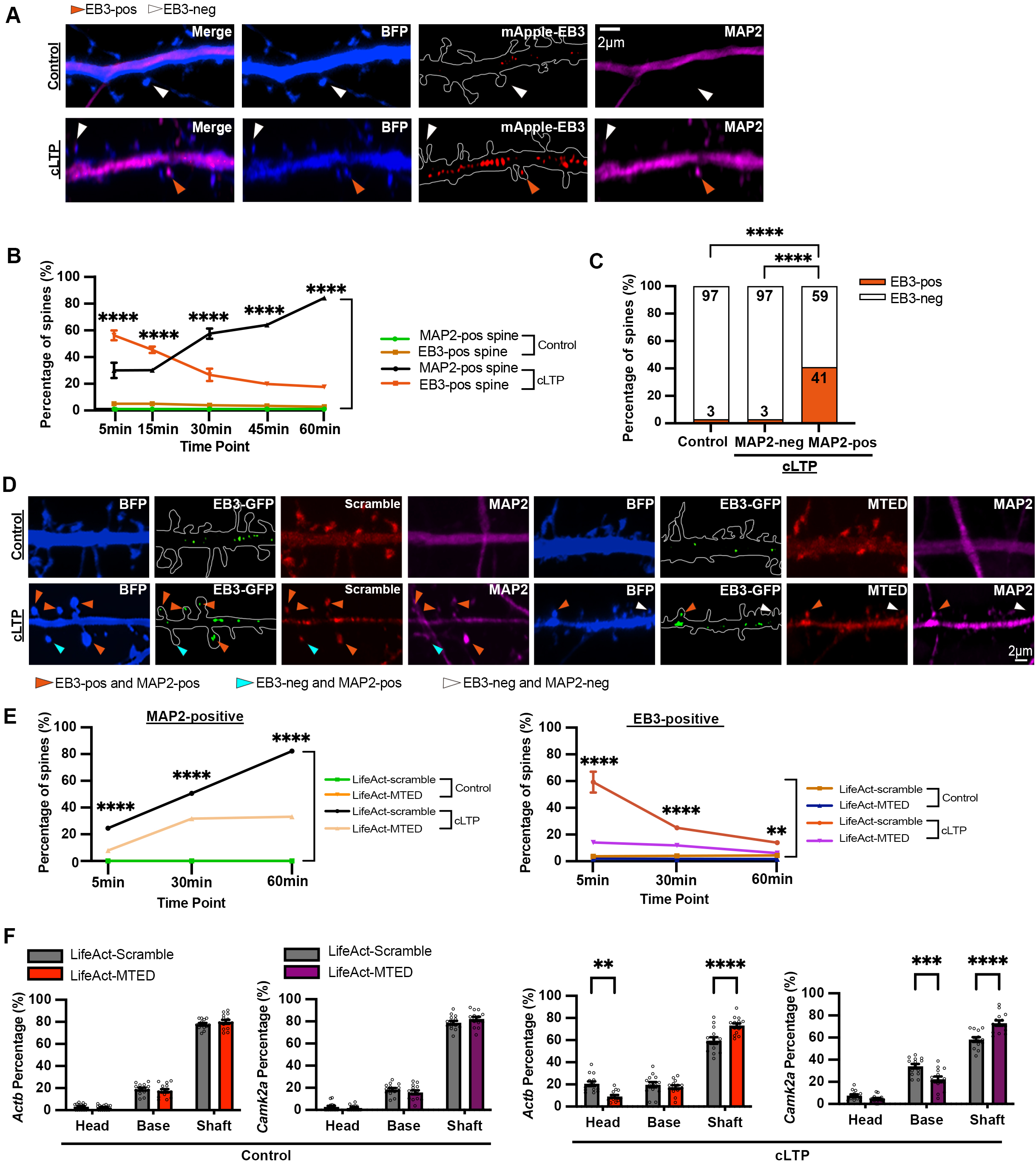
Localization of mRNAs to dendritic spines depends on microtubule entry. (A) EB3 invasion in dendritic spine heads after cLTP. Representative images of dendrites of hippocampal neurons (21 DIV) transfected with EB3-mApple and BFP (13 DIV) and subjected to control treatment or cLTP stimulation. EB3-positive spine is indicated by orange arrowhead; EB3-negative spine is indicated by white arrowhead. (B) Induced EB3 localization at dendritic spine head precedes MAP2 entry in response to cLTP. Rat hippocampal neurons were co-transfected with EB3-mApple and BFP (13 DIV) and subjected to cLTP stimulation at 21 DIV. Neurons were processed at the indicated time points (5, 15, 30, 45, 60 min) after cLTP for immunostaining with BFP and MAP2. The percentages of EB3-positive spines and MAP2-positive spines were quantified at each time point. Results were pooled from two independent experiments; 3,117–4,180 spines from 16 neurons per condition were quantified. Data are mean ± SEM; *\*\*\*\*p < 0.0001*; two-way ANOVA with Šídák’s multiple comparisons test. (C) EB3 granules are specifically enriched in MAP2-positive spines following cLTP. The proportion of EB3-positive spines was quantified in three categories: spines from unstimulated control neurons, MAP2-negative spines after cLTP and MAP2-positive spines after cLTP induction. EB3 showed a preferential association with MAP2-positive spines after synaptic stimulation. Results were pooled from two independent experiments; 884–4,180 spines from 16 neurons per condition were quantified. *\*\*\*\*p < 0.0001*; chi-square test. (D) Inhibition of microtubule polymerization into dendritic spines by LifeAct-MTED disrupts the localization of EB3 and MAP2 in spine heads. Representative images of dendrites of hippocampal neurons transfected with BFP and LifeAct-Scramble or LifeAct-MTED (13 DIV) and subjected to cLTP stimulation at 21 DIV. EB3- and MAP2-positive spines are indicated by orange arrowheads; EB3-negative and MAP2-positive spines are indicated by blue arrowheads, EB3-negative and MAP2-negative spines are indicated by white arrowheads. (E) Specific disruption of microtubule polymerization into dendritic spines by LifeAct-MTED significantly reduced the percentage of spines containing EB3 or MAP2. Results were pooled from two independent experiments; 3,025–5,550 spines from 15-16 neurons per condition were quantified. Data are mean ± SEM; *\*\*p < 0.01*, *\*\*\*\*p < 0.0001*; two-way ANOVA with Šídák’s multiple comparisons test. (F) Disruption of microtubule polymerization into dendritic spines by LifeAct-MTED significantly impairs the cLTP-induced localization of *Actb* and *Camk2a* mRNAs at the spine head and spine base, respectively. Percentage of localization for *Camk2a* and *Actb* mRNA granules in the dendritic shaft, spine base and spine heads in primary basal dendrites across different experimental conditions was quantified. Results were pooled from two independent experiments; 310-468 mRNA puncta from 15 neurons per condition were quantified. Data are mean ± SEM; *\*\*p < 0.01, ****p < 0.0001*; two-way ANOVA with Šídák’s multiple comparisons test.

To test the causal role of microtubule invasion in mRNA localization at dendritic spine, LifeAct-MTED, which couples the Microtubule Elimination Domain (MTED) to the actin-binding peptide LifeAct, was expressed in hippocampal neurons to specifically disrupt microtubule invasion into dendritic spine (Fig. 4D). This tool leverages the high actin concentration in spines to locally eliminate microtubules, thereby inhibiting their invasion without altering overall microtubule dynamics in the dendritic shaft or general neuronal health (Holland et al., 2024). The spine density was relatively unaffected by LifeAct-MTED expression (Fig. S4A). As expected, cLTP triggered a significant increase in EB3 density in the neurons that expressed the LifeAct-scramble control (Fig. S4B). However, expression of LifeAct-MTED profoundly suppressed the influx of both the dynamic microtubule tip protein EB3 and the stabilizing protein MAP2 into spines across all time points after cLTP (Fig. 4E, Fig. S4C), confirming that the localization of both EB3 and MAP2 to spines is strictly dependent on microtubule entry. Most importantly, the blockade of microtubule invasion significantly reduced the localization of *Actb* and *Camk2a* mRNAs to the spine heads and spine base, respectively, after cLTP (Fig. 4F). These findings provide direct evidence that microtubule entry into spines is a prerequisite for the activity-dependent synaptic targeting of *Actb* and *Camk2a* mRNAs.

### Transport of *Actb* and *Camk2a* mRNAs to dendritic spine specifically requires the kinesin motor protein KIF5A but not the closely related KIF5B

Kinesin motor moves towards the plus end of microtubule to mediate anterograde transport in axons and dendrites (Hirokawa and Takemura, 2004; Hirokawa and Tanaka, 2015). The kinesin-1 family, which consists of three homologous members KIF5A, KIF5B and KIF5C, are the major molecular motors associated with mRNA transport in the mouse brain. Given their non-overlapping functions in dendritic spine morphogenesis (Zhao et al., 2020), it is possible that different mRNAs utilize different motor proteins for trafficking. To investigate whether the microtubule-dependent mRNA delivery to dendritic spines is mediated by specific kinesin motors, we focus on the neuron-specific kinesin-1 motor KIF5A. Knockdown of KIF5A specifically reduced the density of *Camk2a* mRNA in apical dendrites, while the *Actb* mRNA density was unaffected (Fig. 5A-B). The somatic fluorescence intensity of *Camk2a,* but not *Actb* mRNA, also decreased upon KIF5A knockdown (Fig. 5C). Nonetheless, the proximo-distal distribution of *Camk2a* mRNA puncta along apical dendrites was significantly disrupted upon KIF5A knockdown (Fig. S4D), confirming that the transport of *Camk2a* mRNA depends on KIF5A. Knockdown of KIF5A also completely abolished the cLTP-induced localization of *Camk2a* at the spine base as well as *Actb* in the spine heads. Notably, the localization of both mRNAs at dendritic spine could only be rescued by the co-expression of RNAi-resistant KIF5A (KIF5Air) but not KIF5B (KIF5Bir) (Fig. 5D-E), indicating that the two similar kinesin motors do not share redundant functions in the synaptic delivery of the mRNAs and KIF5A is specifically required for this process. Neither the spine density, mRNA abundance nor the cLTP-induced spine enlargement was significantly altered by KIF5A knockdown (Fig. S4E-G), suggesting that the impaired mRNA localization at dendritic spine is not due to reduced spine number nor lack of response to the cLTP stimulation. These findings therefore identify KIF5A as the crucial and specific motor that mediates the cLTP-induced and microtubule-based transport of mRNAs to dendritic spines. Importantly, the observation that KIF5A depletion specifically interferes with the cLTP-induced *Actb* localization at dendritic spine (Fig. 5E) without disrupting its overall transport to the dendrites (Fig. 5B, Fig. S4D) reveals that the two types of mRNA transport (long-range soma-to-dendrite and the short-range delivery to dendritic spine) can be uncoupled and independently carried out by different motor proteins.

**Figure 5.**
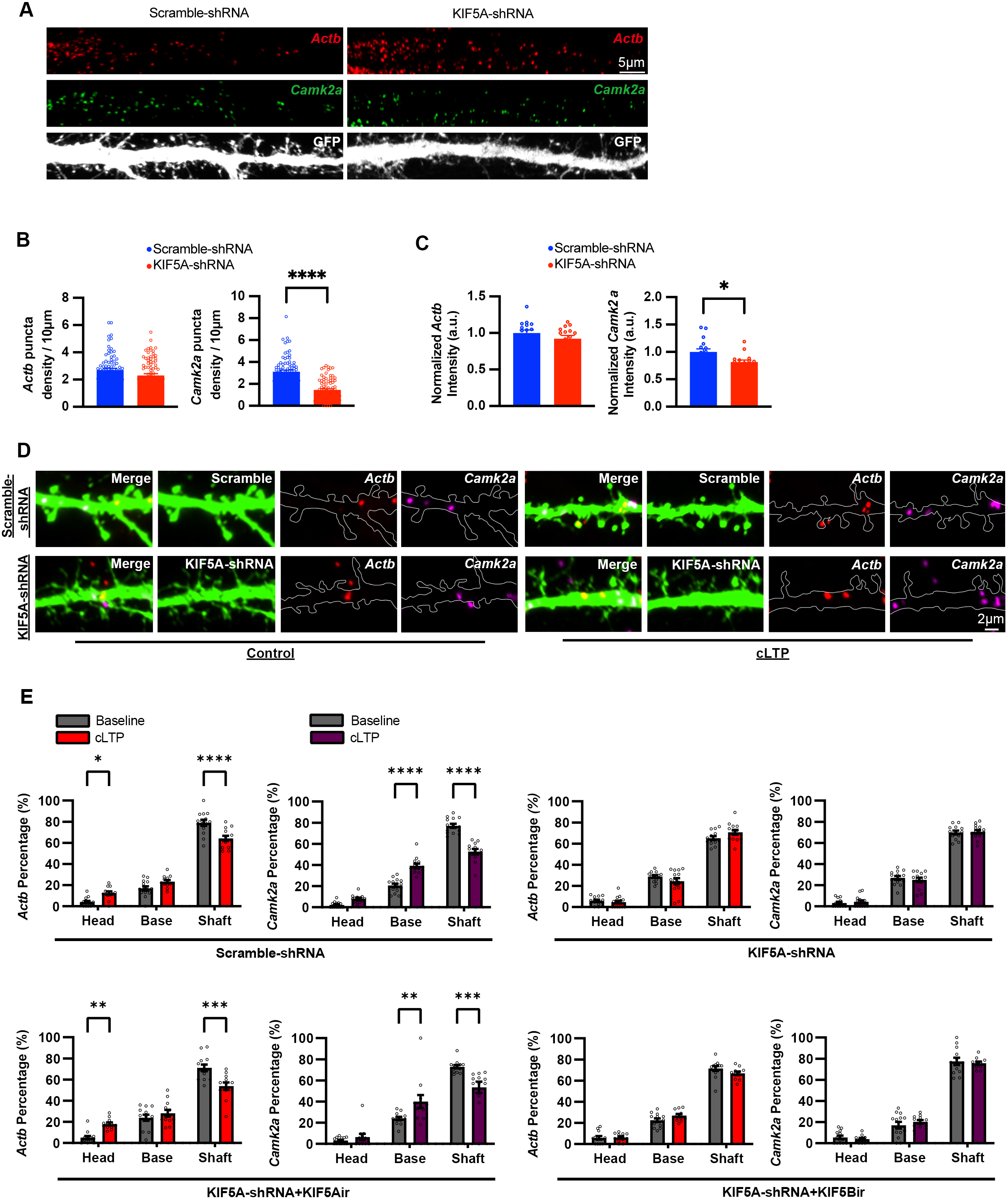
The kinesin motor protein KIF5A is specifically required for the transport of *Actb* and *Camk2a* mRNAs to dendritic spines after cLTP. (A) Representative images of apical dendrites of rat hippocampal neurons (21 DIV) transfected with either GFP-scramble-shRNA or GFP-KIF5A-shRNA (13 DIV). The Actb and Camk2a mRNAs were visualized by dual-color smFISH. (B) KIF5A knockdown specifically reduces the density of *Camk2a,* but not *Actb,* mRNA puncta in apical dendrites. Each data point represents one dendrite. Results were pooled from three independent experiments, 25-26 neurons per condition were quantified. Data are mean ± SEM, *\*\*\*\*p < 0.0001*, Student’s t-test. (C) KIF5A knockdown specifically reduces the level of *Camk2a*, but not *Actb,* in the cell body, as indicated by the average intensity of the mRNA signals. Each data point represented one neuron. Results were pooled from two independent experiments; 25-26 neurons per condition were quantified. Data are mean ± SEM, *p < 0.05, Student’s t-test. (D) Representative images of primary basal dendrites of rat hippocampal neurons transfected with GFP-scramble-shRNA or GFP-KIF5A-shRNA (13 DIV), followed by cLTP stimulation at 21 DIV. Neurons were fixed for dual-color smFISH 60 min after cLTP to visualize *Camk2a* (magenta) and *Actb* (red). (E) KIF5A but not the closely related homolog KIF5B is required for the cLTP-induced *Actb* and *Camk2a* localization at dendritic spine. Percentage of *Camk2a* (magenta) and *Actb* (red) localization in the dendritic shaft, the spine base and the spine head of primary basal dendrites across the experimental groups: scramble shRNA, KIF5A shRNA, KIF5A shRNA + KIF5Air, and KIF5A shRNA + KIF5Bir was quantified under control and cLTP conditions. Results were pooled from two experiments; 128-304 mRNA puncta from 11-14 neurons per condition were quantified. Data are mean ± SEM; *\*p < 0.05, **p < 0.01, ***p < 0.001, ****p < 0.0001*; two-way ANOVA with Šídák’s multiple comparisons test.

The C-terminals of KIF5A (residues 808–1027) and KIF5B (residues 808–963) contain the cargo-binding domain and the diverse carboxyl-tail that determines their functional specificity in dendritic spine morphogenesis (Zhao et al., 2020). We therefore hypothesize that the C-terminal domains of the three different KIF5s might associate with distinct sets of RBPs in neurons that could contribute to RNA transport specificity. To explore this possibility, we perform APEX2-mediated proximity labelling to identify putative proteins that selectively interact with the different KIF5 motors in neuronal context. Fusion proteins comprising APEX2 and the C-terminal regions of KIF5A, KIF5B, or KIF5C were constructed and electroporated into primary cortical neurons, with the plasmid containing only the APEX2 backbone as negative control (Fig. 6A-C). Biotinylated proteins were enriched by streptavidin beads and identified by mass spectrometry. Three biological replicates were included for each experimental condition, and the PCA plot showed strong consistency among the triplicates (Fig. 6D). 500 proteins were identified to be enriched with statistical significance in the KIF5-APEX2 proteome relative to the APEX2 control (Fig. 6E). Several RBPs reported to be localized in the dendrites, including KHDRBS1 (also known as SAM68), were enriched by all three KIF5s. Notably, we also identified dendritic RBPs that showed selective enrichments by specific KIF5 motors. For example, CAPRIN1 and YBX1 were enriched only by KIF5B and KIF5C, whereas FMRP was enriched selectively by KIF5A (Fig. 6F-G). These findings suggest that the three different KIF5 motors could indeed couple to different RBPs that may constitute a potential mechanism for the heterogeneous transport of mRNAs.

**Figure 6.**
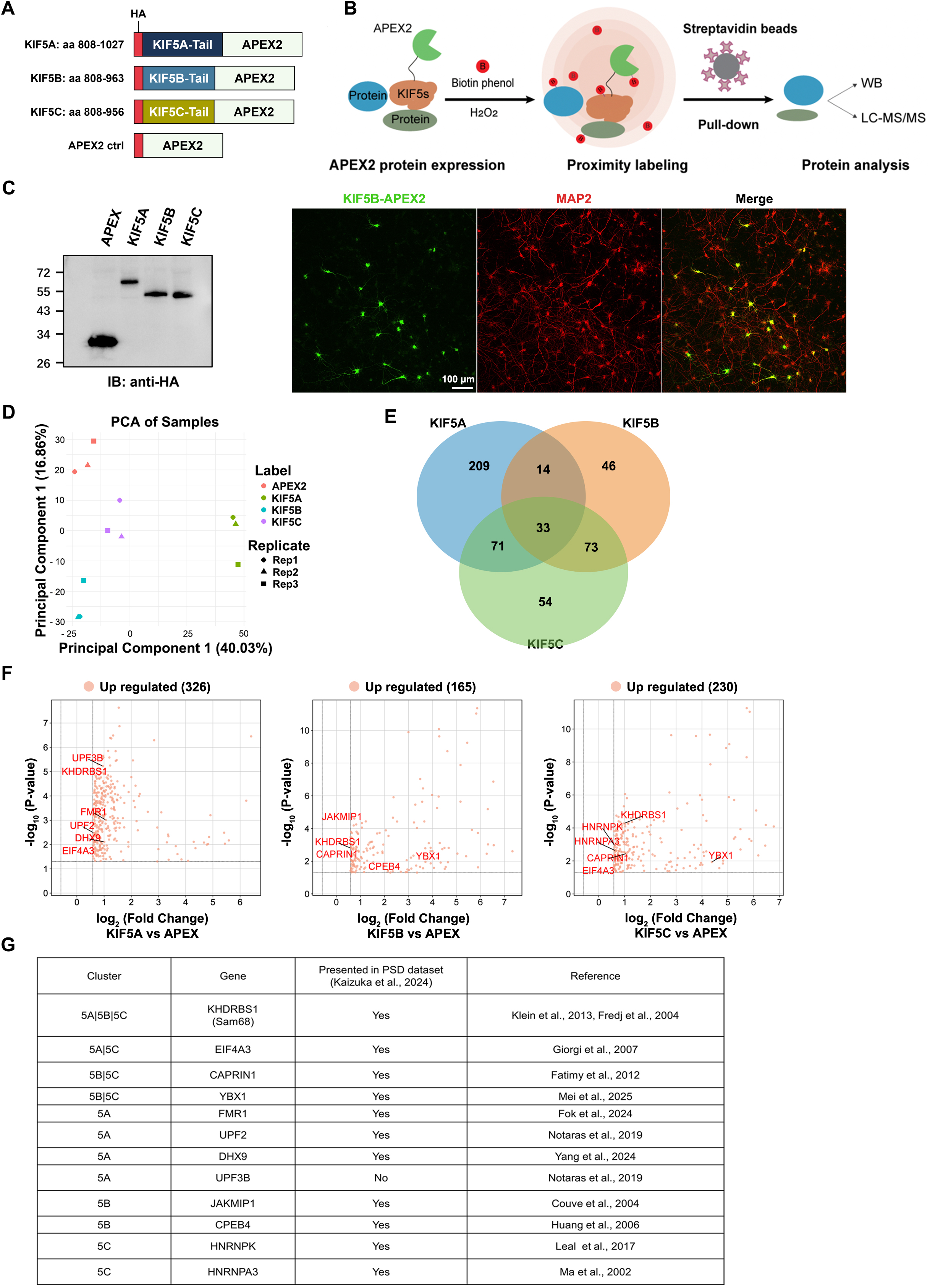
APEX-mediated proximity labelling proteomics identifies dendritic RBPs near the different KIF5 homologs in neurons. (A) Schematic representation of the KIF5-APEX2 constructs. The C-terminal half of mouse KIF5A (808-1027 aa), KIF5B (808-963 aa) and KIF5C (808-956 aa) were fused to APEX2, which also contained a HA tag at the N-terminus. (B) Schematic representation of the KIF5-APEX2 proximity labeling method. APEX2 oxidizes biotin-phenols to biotin-phenoxyl radicals in the presence of H_2_O_2_. These radicals subsequently biotinylate nearby proteins within a range of < 20 nm. The biotinylated proteins are enriched using streptavidin beads and applied to subsequent analyses such as Western blotting and mass spectrometry. (C) Expression of the various KIF5-APEX2 fusion proteins in cultured cortical neurons. The HA-tagged KIF5-APEX2 plasmids were electroporated into primary cortical neurons at 0 DIV and were harvested at 14 DIV for Western blot analysis (left panel) or immunofluorescence staining (right panel) using anti-HA and MAP2 antibodies. (D) Principal component analysis (PCA) plot of the KIF5-APEX2 mass spectrometry results. Each data point corresponds to one MS sample. The plot demonstrates a clear separation between the four experimental groups. (E) Venn diagram showing the overlap between the KIF5A, KIF5B, and KIF5C proximity-proteomes. Numbers in the diagram represent the number of proteins that are significantly enriched (adjusted *p*-value < 0.05, and fold change > 1.5 as compared to the APEX2-backbone negative control). (F) Volcano plots showing the RBPs enriched in the proximity of KIF5A, KIF5B, and KIF5C. Significantly enriched proteins (adjusted *p*-value < 0.05 and fold change > 1.5) are colored in red. Several known RBPs are annotated, revealing their association with specific KIF5 homologs. (G) Table summarizing the various dendritic RBPs enriched in the different KIF5-proximity proteomes. Previous studies demonstrating the dendritic localization of these RBPs were listed in the last column.

### *Actb* and *Camk2a* mRNAs are localized in proximity to different synaptic proteins in neurons

To further confirm the heterogeneous localization of *Actb* and *Camk2a* at dendritic spines and identify potential proteins that underlie their differential targeting, we attempt to characterize the proteins that are situated near the two mRNAs in their native cellular context via the proximity labelling approach called CRISPR-Assisted RNA-Protein Interaction Detection (CARPID). Unlike most RNA-protein proximity-labeling methods that start with a protein of interest as the target, this CRISPR-based strategy is RNA-centric and involves biotin-labelling of proteins located within ∼20 nm to the RNA of interest (Yi et al., 2020). Towards this end, cortical neurons which were used instead of hippocampal neurons because of the large quantity of proteins needed as the starting material, were transfected with two constructs: one expressing the dCas13x.1 fused with the biotin-ligase BASU and the other expressing a set of three gRNAs targeting either *Actb* or *Camk2a* mRNA, respectively (Fig. 7A-B). A scrambled gRNA was used as the negative control in parallel. The electroporation efficiency of CARPID plasmids was verified by immunostaining (Fig. S5A). Each experimental condition contained biological triplicates which showed strong consistency (Fig. 7C, Fig. S5B). The patterns of enriched proteins in the two mRNA clusters exhibited clear differences (Fig. 7C-D). Of the 521 proteins enriched by *Camk2a* mRNA, 361 of them were enriched only in the *Camk2a* but not the *Actb* cluster; whereas among the 464 total enriched by *Actb* mRNA, 304 proteins were enriched specifically in the *Actb* mRNA cluster (Fig. 7E, Supplementary Table 1). GO analysis revealed that the two clusters of proteins are involved in different biological processes and cellular compartments. (Fig. 7F). Of note, ZBP1 (IGF2BP1) and Sam68 (KHDRBS1) which are known RBPs that transport *Actb* mRNA in neurons (Tiruchinapalli et al., 2003; Klein et al., 2013), were successfully enriched in the *Actb* cluster by CARPID, thereby validating the method in isolating RNA-interacting proteins from neurons. To further verify the CARPID method, we employed cartRAPID, an algorithm that estimates RNA-protein binding propensity based on physicochemical properties of the molecules (Bellucci et al., 2011), to retrieve proteins predicted to associate with *Actb* and/or *Camk2a* mRNAs. Alignment of these proteins with our CARPID list identified overlapping candidates (Supplementary Table 2), suggesting that CARPID can identify specific interacting proteins for the mRNAs of interest. To focus on postsynaptic proteins, we first compared our CARPID data with the published meta-analyses of PSD proteome datasets encompassing a total of 5,869 proteins (Kaizuka et al., 2024). Among these postsynaptic proteins, 290 were specifically enriched in the *Camk2a* mRNA cluster and 185 were enriched in the *Actb* one (Fig. 7G). Next, we specifically selected the GO categories that include the term “postsynaptic”. Surprisingly, there was a striking difference between the proteins enriched by the two mRNA clusters, as these “postsynaptic” GO terms only came up in the *Camk2a*-proximity proteome (Fig. 8A).

**Figure 7.**
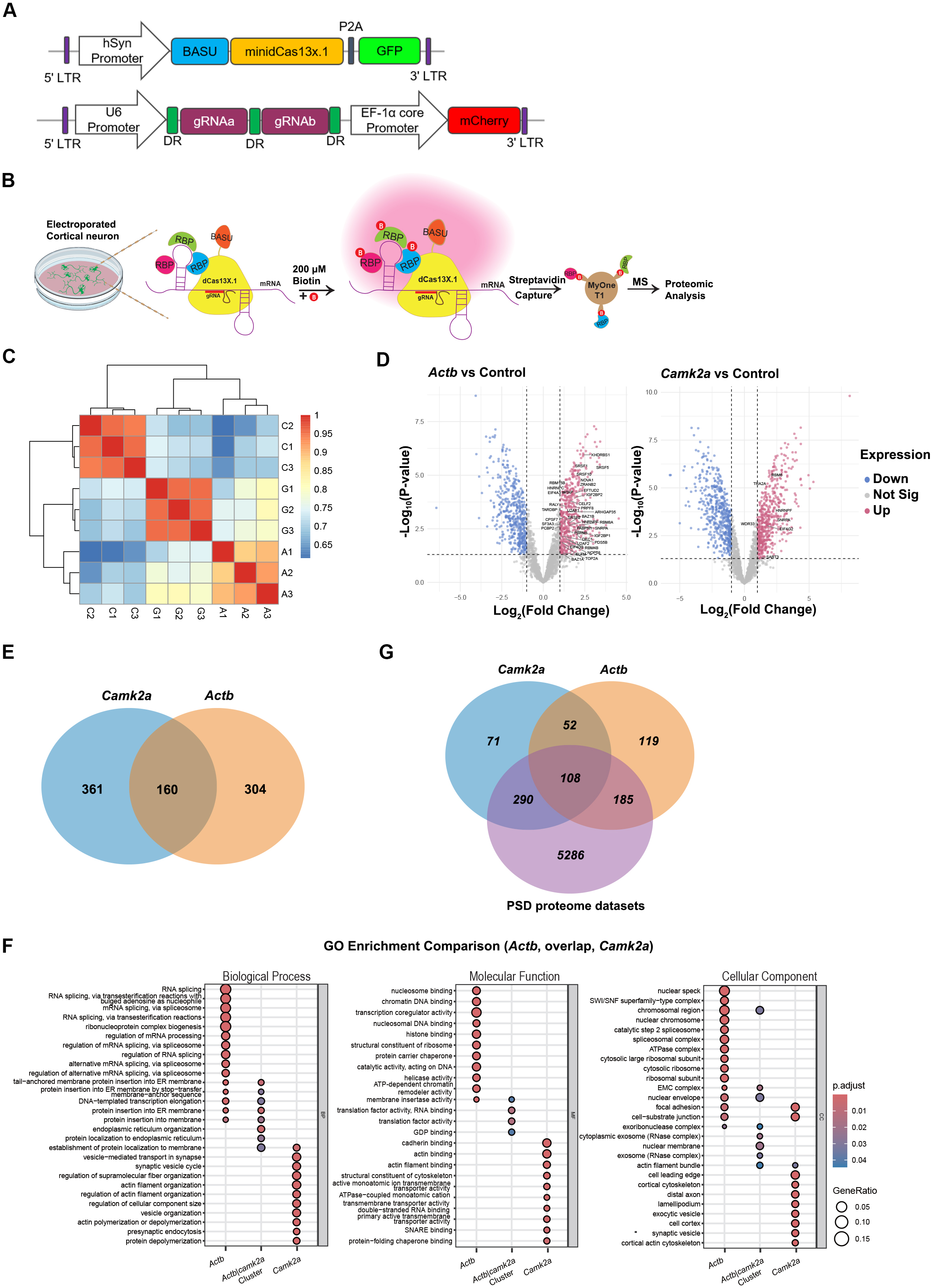
CARPID proximity labelling identifies different proteins enriched by *Actb* and *Camk2a* mRNAs in neurons. (A) Schematic representation of the CARPID plasmids. The minidCas13X.1 were tagged with the biotin ligase BASU and GFP. Two gRNA sequences targeting two adjacent loci on either the *Actb* or *Camk2a* transcript were tagged with mCherry. Three different gRNA plasmids that target different regions of each mRNA were designed and transfected together into cortical neurons. (B) Overview of the CARPID-MS workflow. Cortical neurons were electroporated with the CARPID plasmids and treated with biotin (200 μM) at DIV16 (biotin is denoted by the red circles labeled ‘B’). Protein interactors within the proximity of the bait mRNAs were labelled. After streptavidin pulldown, samples were subjected to LC-MS/MS. (C) Correlation Matrix heat map shows the Pearson correlation coefficients (r) between *Camk2a*, *Actb*, and control gRNA. Cells were shaded according to the strength of relationships based on the Pearson correlation coefficients (r) (p < 0.001). (D) Volcano plots illustrate the enrichment of proteins (up, pink dots) by *Actb* gRNAs (left) and *Camk2a* gRNAs (right), as compared to the control gRNA (Fold Change ≥ 2, P value < 0.05). RNA-binding proteins (RBPs) were annotated. (E) Venn diagram shows the common and unique proteins between the *Camk2a*-proximity (blue) and *Actb*-proximity proteomes (orange). A total of 361 proteins were identified exclusively in the *Camk2a*-proximity proteome, and 304 proteins were identified exclusively in the *Actb*-proximity proteome. 160 proteins enriched by both mRNAs were identified. (F) Gene Ontology (GO) terms enriched among the *Actb*- and *Camk2a*-proximity proteins as well as the overlapping proteins. Gene enrichment percentages are indicated by sizes of the dots. Dot color gradients represent Benjamini–Hochberg adjusted *p*-values (*p* < 0.05). (G) Venn diagram that compares the CARPID data with the PSD proteome datasets. The overlap between proteomes enriched by the two mRNA clusters determined by CARPID and the previously published PSD proteome database (purple) encompassing a total of 5,869 proteins is shown.

**Figure 8.**
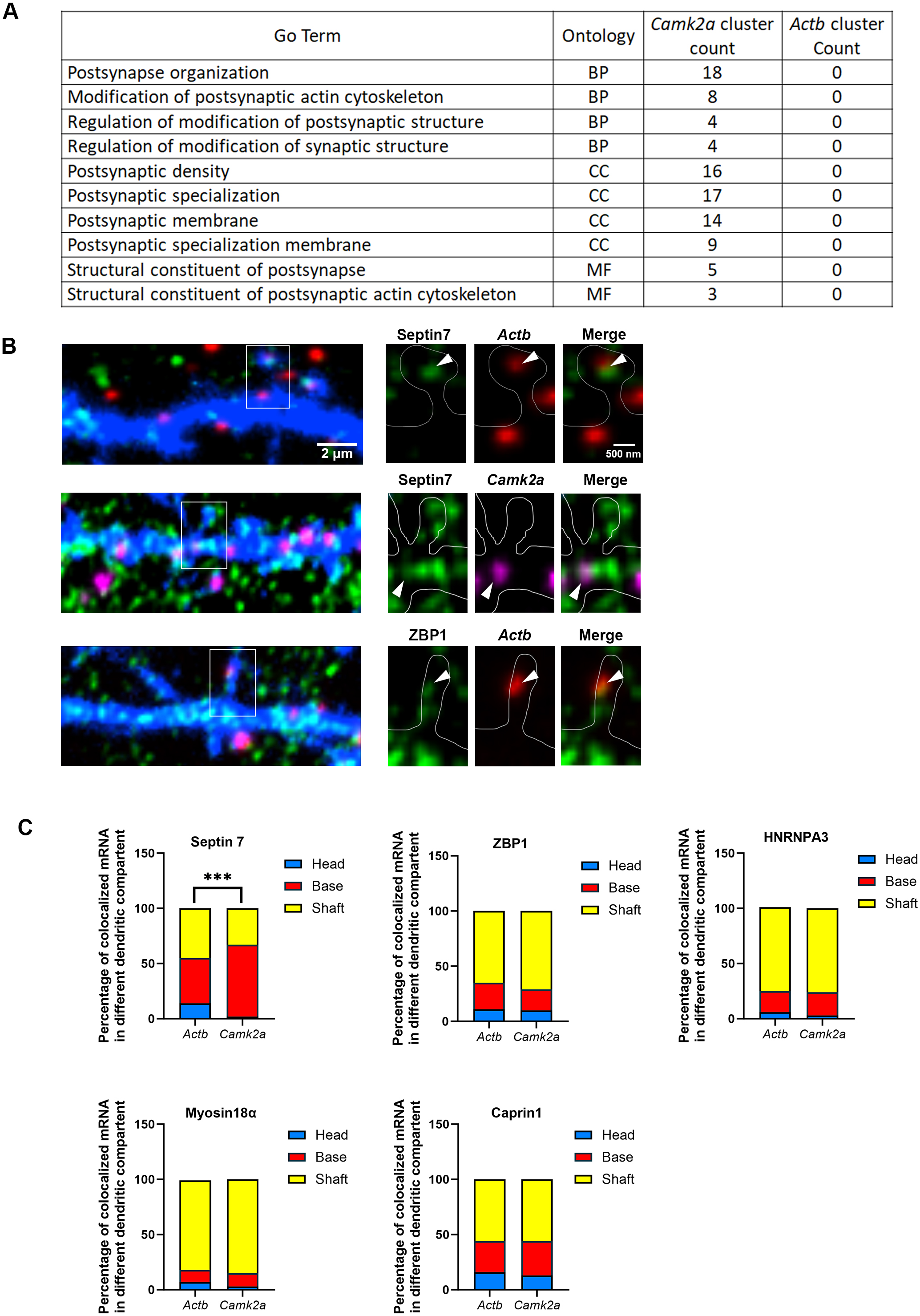
Different dendritic distribution of the co-localization between *Actb*/*Camk2a* mRNAs and the proteins identified by CARPID-MS in neurons. (A) Table summarizing the number of proteins from the *Camk2a*- and *Actb*-proximity proteomes that are present in the GO terms related to postsynaptic functions. Only *Camk2a*-proximity proteins were identified in these GO terms. (B) Representative images of dendritic segments of rat hippocampal neurons (21 DIV) transfected with BFP (13 DIV). Enlarged views showing the co-localized RNA-protein at the base (Septin7-*Camk2a*) or head (Septin7-*Actb* and ZBP1-*Actb*) of dendritic spines. (C) Majority of Septin7-mRNA co-localization was detected at the base of dendritic spines, and significant difference in distribution was observed between Septin7-*Actb* and Septin7-*Camk2a* co-localized puncta at the different dendritic compartments. The percentages of *Actb* and *Camk2a* mRNA puncta colocalized with the different proteins at three dendritic compartments: the dendritic shaft (yellow), the spine base (red), and the spine head (blue) were shown. Results were pooled from three independent experiments; 15-27 neurons were imaged and 232-309 mRNA puncta from 32-65 dendrites were analyzed. *\*\*\*p < 0.001*, chi-square test.

Next, five proteins (ZBP1, Septin7, hnRNPA3, Myo18a and CAPRIN1) from the CARPID screen were selected for immunostaining and dual-color smFISH in hippocampal neurons to examine their co-localization with *Actb* and *Camk2a* at different subdomains of dendritic spines within the same neurons (Fig. 8B). Co-localization of the two mRNAs, ranging from 12% to 36%, were observed with all five selected proteins on the dendrites, hence confirming the CARPID proximity labeling results. We then categorize distribution of the co-localized mRNA-protein puncta across the three dendritic compartments (spine head, spine base and dendritic shaft). The two mRNAs co-localize with the four proteins (ZBP1, hnRNPA3, Myo18a and CAPRIN1) with comparable distribution patterns, and most of the co-localized puncta were present on the dendritic shaft (Fig. 8C). In contrast, a distinct distribution pattern of mRNA co-localization was observed for Septin7, a cytoskeletal protein that promotes crosstalk between actin and microtubule filaments (Nakos et al., 2022) and is enriched at the base of dendritic spines (Tada et al., 2007; Xie et al., 2007). First, compared to the other four proteins, much higher Septin7-mRNA co-localization was observed at the spine base. Second, the distribution was significantly different between the two mRNAs (Chi-square test, p-value 0.0003), with majority (∼64%) of the *Camk2a*-Septin7 co-localization detected at the spine base and rarely (2%) found in the spine head; whereas considerable proportion (∼14%) of Septin7-*Actb* co-localized puncta was detected in the spine head (Fig. 8C). These findings suggest that Septin7, which acts as a diffusion barrier and controls material transport between the spine head and the dendritic shaft (Ewers et al., 2014), might be involved in the docking of mRNAs at the spine base, as well as contribute to the specific distribution of *Actb* and *Camk2a* within dendritic spines. From a broader perspective, our study demonstrates CARPID as a useful method to identify the protein interactomes of a particular mRNA of interest at various subcellular compartments of neurons, including the different subdomains of dendritic spine.

## DISCUSSION

While the transport of mRNAs along dendrites is well-documented, whether and how different transcripts are differentially targeted to and distributed within individual dendritic spines remains unclear. By simultaneously visualizing *Actb* and *Camk2a* in the same neurons for head-to-head comparison, our study reveals a high degree of spatial heterogeneity in the sub-spine localization of the two dendritic mRNAs. Although both mRNAs are more frequently detected at the base than within the head of dendritic spines and cLTP promotes the transport of both transcripts to dendritic spines, differential distribution is observed with cLTP specifically induces *Actb* localization to the spine head, while the induced targeting of *Camk2a* occurs mostly at the spine base. Another aspect of specificity is demonstrated by the stronger preference of *Camk2a* to the head of MAP2-positive and synaptopodin-enriched spines when compared to *Actb*. Our findings further indicate the invasion of microtubules and MAP2 as the prerequisite “gating” mechanism for this activity-dependent mRNA recruitment that also depends on the microtubule-based kinesin motor KIF5A. Finally, CARPID proximity-labeling-MS identify distinct protein interactomes of these mRNAs, including postsynaptic proteins. From the proteomics, we uncover the cytoskeletal protein Septin7 being co-localized with *Actb* and *Camk2a* at different sub-domains of dendritic spines. Collectively, our findings suggest a multi-step model where cytoskeletal dynamics and interactions with specific proteins orchestrate the precise sub-compartmentalization of mRNAs to support synapse-specific plasticity.

### The Spine Base as a Strategic Reservoir of mRNAs

Based on the increased co-localization of *Actb* and *Camk2a* at the spine base, we suggest this dendritic subdomain as the primary docking site for the precise sub-synaptic organization that mirrors the localization of *Arc* (Dynes & Steward, 2012). We propose that the spine base may function as a strategic “hot spot,” potentially optimizing the trade-off between proximity and spatial constraints. Structurally, the spine head is a highly restricted compartment with densely packed scaffold proteins and receptors in the PSD (Swulius et al., 2010). Anchoring mRNAs at the spine base, which is enriched with translation machinery and polyribosomes, likely allows neurons to maintain a pool of “poised” transcripts close enough to the synapses but without overcrowding the spine head, thereby leaving the PSD to carry out the important function of neurotransmission. In addition, we postulate that this mRNA reservoir at the spine base provides functional flexibility. Instead of being “locked” within the head of a particular spine, localization at the base facilitates the redistribution of mRNAs to neighboring synapses if local activity does not reach the threshold for plasticity. Our time-lapse imaging shows that *Actb* granules that dock underneath dendritic spines are still dynamic and can soon move away from the initial anchoring spines, hence supporting the notion that docking at the spine base serves as a checkpoint rather than a terminal destination. Upon synaptic stimulation, the reservoir underneath the spine base might become mobilized, which is supported by our observed reduction of *Actb*-*Camk2a* co-localization at the spine base after cLTP. The subsequent translocation of *Actb* to the spine head conceivably ensures that the synthesis of β-actin is spatially confined to the active spine, providing the structural substrate required for spine enlargement while minimizing effect at unstimulated sites (Yoon et al., 2016).

### Activity-Dependent Sorting into Distinct Spine Subdomains and Populations

Our study reveals that cLTP induction leads to divergence in the spatial organization of *Actb* and *Camk2a* mRNAs. We postulate that this spatial segregation likely reflects the distinct properties and functional demands of their encoded proteins during plasticity. β-actin is the primary building block for spine enlargement. Upon polymerization into F-actin, the large size of the polymer makes it difficult to translocate elsewhere. Therefore, the specific localization of *Actb* mRNAs within the spine head is critical to concentrate the synthesis of structural components directly at the site of remodeling (Nakahata and Yasuda, 2018; Yoon et al., 2016). In contrast, the retention of *Camk2a* mRNA at the spine base might fulfil a different regulatory logic. Because of its small size and diffusible nature, CaMKIIα locally synthesized at the spine base would readily move into the spine head. Given the abundance of CaMKIIα in the PSD, it is conceivable that *Camk2a* mRNA translation needs to be highly effective. As such, it might be desirable to mainly synthesize CaMKIIα at the spine base where polyribosomes and other translation machinery are likely to be more abundant compared to the spine head.

Despite their preferred targeting to the spine base after cLTP, we do observe *Camk2a* within the spine heads. In this regard, it is interesting that, compared to *Actb* within the same neurons, these *Camk2a* granules in the spine heads exhibit a higher degree of selectivity towards the MAP2-positive and SP-enriched spines. In support of the latter, we found from our CARPID data the *Camk2a*-proximity proteins show larger overlap with the spine apparatus proteome (12 out of 140 spine apparatus proteins from Falahati et al., 2022) than the *Actb*-proximity ones (only 2 out of 140). These findings suggest that specific transcripts such as *Camk2a* are localized in proximity to the spine apparatus. MAP2 invasion is a hallmark of structural plasticity, often marking spines that have been functionally potentiated (Kim et al., 2020); whereas SP is a key component of the spine apparatus, an organelle associated with internal calcium stores and local protein synthesis machinery (Konietzny et al., 2023; Pierce et al., 2000; Vlachos et al., 2012). This specific accumulation suggests that *Camk2a* mRNA is not randomly distributed; instead, it is targeted based on the “calcium-handling” capacity of the spine. Since CaMKIIα is a calcium/calmodulin-dependent kinase, positioning its mRNA in SP-enriched spines could be advantageous as the translated product is spatially juxtaposed with internal calcium release sites and potentially facilitating rapid local activation. Conversely, *Actb* mRNA displays a broader distribution, localizing to both MAP2-positive and MAP2-negative spines. This observation implies that while *Camk2a* translation might be prioritized in the more mature or potentiated spines equipped with the spine apparatus, the requirement for actin remodeling is likely more universal, since those dendritic spines that have not yet undergone extensive maturation marked by MAP2/SP accumulation still require continuous actin turnover to maintain their shape or undergo structural changes. Thus, the differential targeting of these two mRNAs might underscore a sophisticated routing mechanism where transcripts are delivered to specific sub-populations of spines based on their local demands.

The CARPID proximity labeling mass spectrometry uncovers that the *Actb* and *Camk2a* mRNAs in neurons enrich distinct sets of proteins in the PSD proteome database, thereby providing additional evidence to support the heterogeneous distribution of mRNAs within specific subdomains of dendritic spine. One of the most intriguing findings arise from the proximity-labeling proteomics is that many GO terms related to postsynaptic membrane and postsynaptic density are only detected in the *Camk2a*-enriched proteins. This suggests that for those *Camk2a* mRNAs located within the spine head, they might be locally synthesized in proximity (probably ∼ 20 nm) to the PSD and synaptic membrane. This might have functional relevance since many PSD proteins are substrates of CaMKIIα. On the other hand, due to the potential overcrowding problem at the PSD or near the postsynaptic membrane, *Actb* whose translation into β-actin is not necessary to be synthesized closely at those sites.

### Microtubule Invasion and KIF5A-Mediated Synaptic Delivery

The actin/myosin motor system is conventionally regarded as the major short-range transport system in neurons (Hammer & Sellers, 2012). Our findings that mRNA localization at dendritic spine is impaired either upon selective inhibition of microtubule polymerization at the spine head or knockdown of the kinesin motor KIF5A suggests that the actin/myosin network alone is insufficient for the synaptic recruitment of mRNAs. We demonstrate that the transient entry of EB3 into spine heads, which labels the dynamic microtubules, precedes the accumulation of MAP2. This could create a temporary “highway” bridging the dendritic shaft and the spine compartment (Miryala et al., 2022). This mechanism potentially explains the preferential targeting of mRNAs to the MAP2-positive spines: since these spines have successfully accommodated microtubules, they have established the necessary infrastructure for cargo delivery. From this point of view, microtubule invasion might function as a checkpoint to ensure that the energy-intensive local translation is deployed only to synapses undergoing active structural plasticity or consolidation (Holland et al., 2024).

Navigating this microtubule network requires specific motor proteins. Our investigation into the Kinesin-1 family reveals a sophisticated division of labor between long-range trafficking and local synaptic capture. For long-range transport from the soma to the dendrite, *Camk2a* and *Actb* mRNAs exhibit distinct motor dependencies. KIF5A knockdown significantly reduces the dendritic density of *Camk2a* but not *Actb*, implying that neurons utilize distinct kinesin homologs to sort mRNAs into different transport granules, potentially allowing for the differential regulation of their abundance in dendrites (Swarnkar et al., 2021). However, despite the redundancy in long-range transport, in which *Actb* can presumably rely on other kinesins for the transport to dendrite upon KIF5A depletion, our knockdown and rescue experiments indicate that KIF5A is indispensable for the cLTP-induced *Actb* localization to the spine heads and cannot be replaced by KIF5B. The observation that KIF5A knockdown prevents *Actb* from entering the spine head, even when the transcript localization on the dendrite is unaffected, demonstrates a mechanistic uncoupling between dendritic transport and synaptic delivery. Consistent with our findings, a very recent study also reports KIF5A functions to transport mRNAs that encode ribosomal proteins to the synapses of human induced pluripotent stem cell (hiPSC)-derived motor neurons (Le et al., 2025). We propose that KIF5A plays a specialized role in the short-range delivery of mRNAs, possibly by transferring RNP granules from the stable microtubules of the dendritic shaft onto the dynamic, invading microtubules in the spine. Alternatively, KIF5A may be unique among Kinesin-1 family members in its ability to navigate the steric constraints of the spine neck or interact with specific guidance factors during cLTP. This requirement, first microtubule invasion as the track followed by its stabilization by MAP2 and finally KIF5A as the motor, provides a multi-step filter to achieve the spatiotemporal control of synaptic mRNA delivery. Nonetheless, we do not rule out the importance of actin filaments for the mRNA targeting to dendritic spine. The entry of microtubules to dendritic spine depends on actin remodeling at the spine base (Schätzle et al., 2018), hence the two types of cytoskeletons likely work together at dendritic spine to achieve the ultimate mRNA delivery. In this regard, it is noteworthy that Septin7, a GTP-binding cytoskeletal protein that facilitates crosstalk of actin and microtubule filaments and regulates kinesin movement (Nakos et al., 2022; Suber et al., 2023), largely localize with both *Actb* and *Camk2a* at the spine base but selectively co-localized with *Actb* within the spine head. Given Septin7 function as a diffusion barrier at the spine base, it is a plausible candidate protein that restricts mRNA mobility at the spine base to facilitate their subsequent local translation. In addition, Septin7 might coordinate the invading microtubules with the actin filaments for specifically bringing the *Actb* granules to the spine head. It will be of great interest to delineate in the future whether and how the oligomeric Septin complex participates in sorting different mRNAs into distinct sub-domains of dendritic spine.

## ACKNOWLEDGEMENTS

We are grateful to Marina Mikhaylova (The Humboldt-Universität zu Berlin) and Yi-Ping Hsueh (Academia Sinica) for the gift of the GFP-synaptopodin plasmid, and Wenjun Xiong (City University of Hong Kong) for the help on electroporation. We thank Angelo Flores and Yutong Wang for assisting the quantification of images, and Xuejun Li for assisting the live imaging of the MS2 reporter mRNA. This study was supported in part by the Research Grant Council of Hong Kong [Collaborative Research Fund (CRF) C1024-22G and General Research Fund (GRF) 11102422 and 11101624]; and the US National Academy of Medicine (NAM) [Healthy Longevity Catalyst Award (Hong Kong) HLCA/M-106/25].

## MATERIALS AND METHODS

**Table 1.**
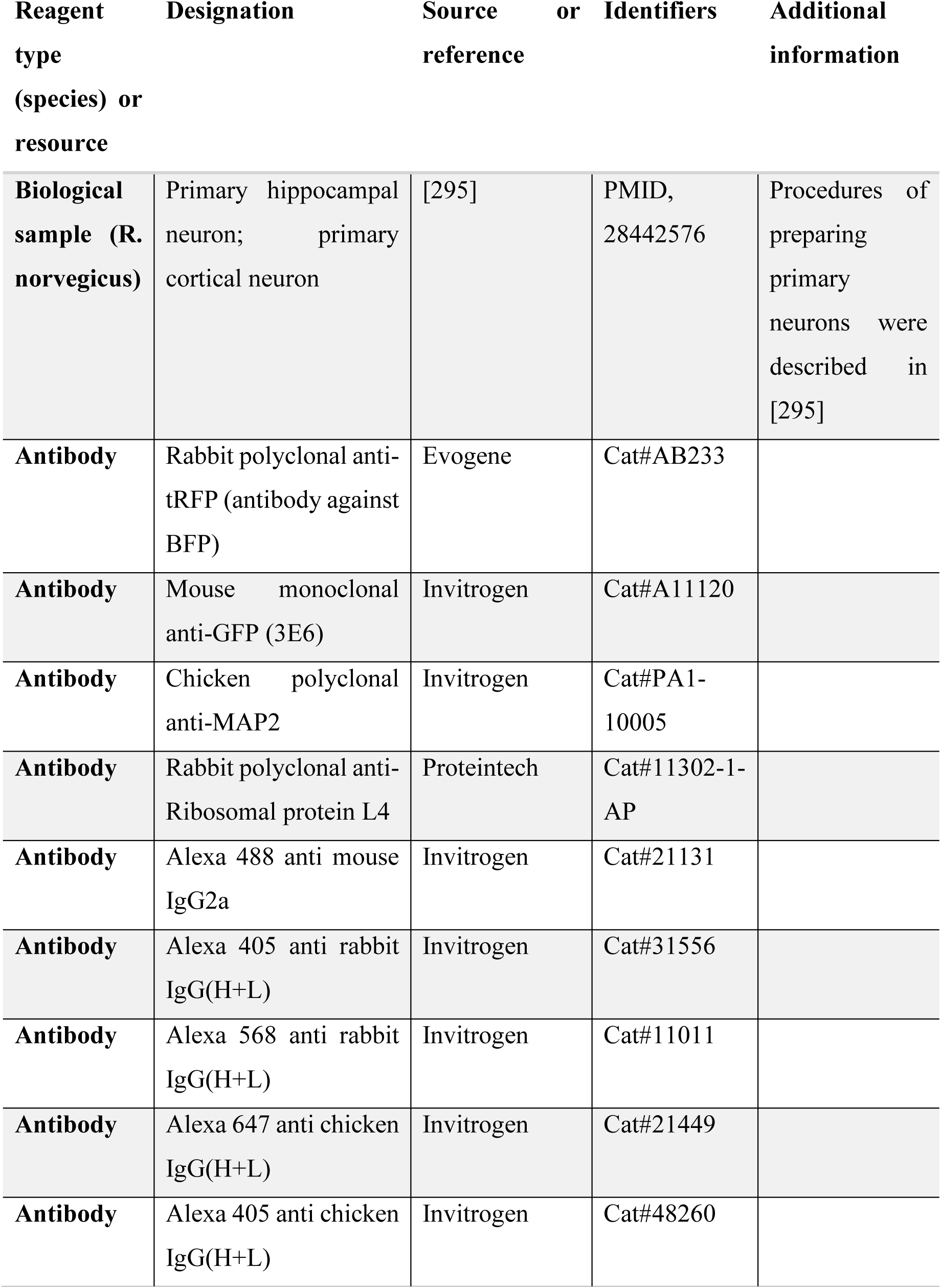
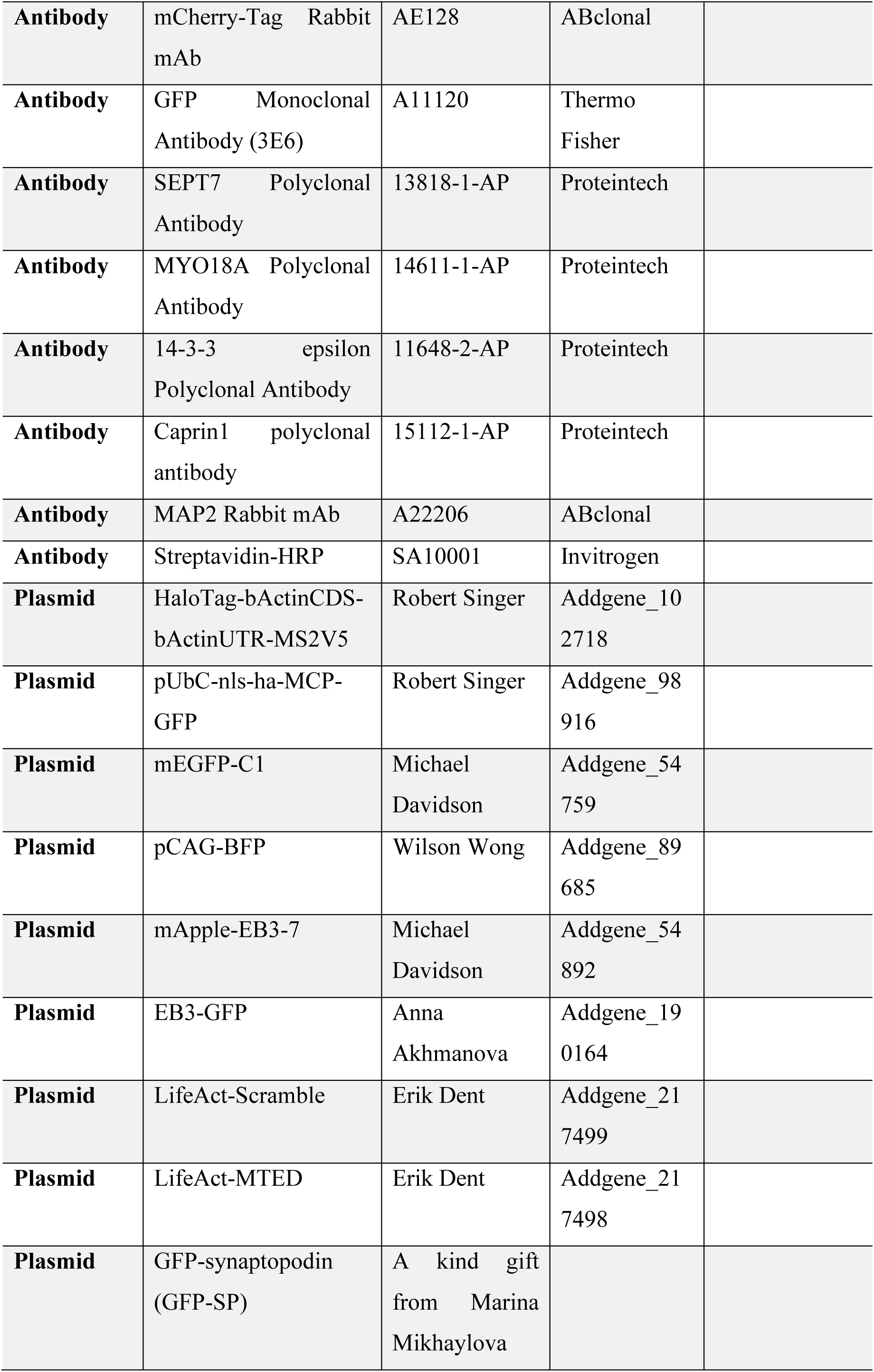
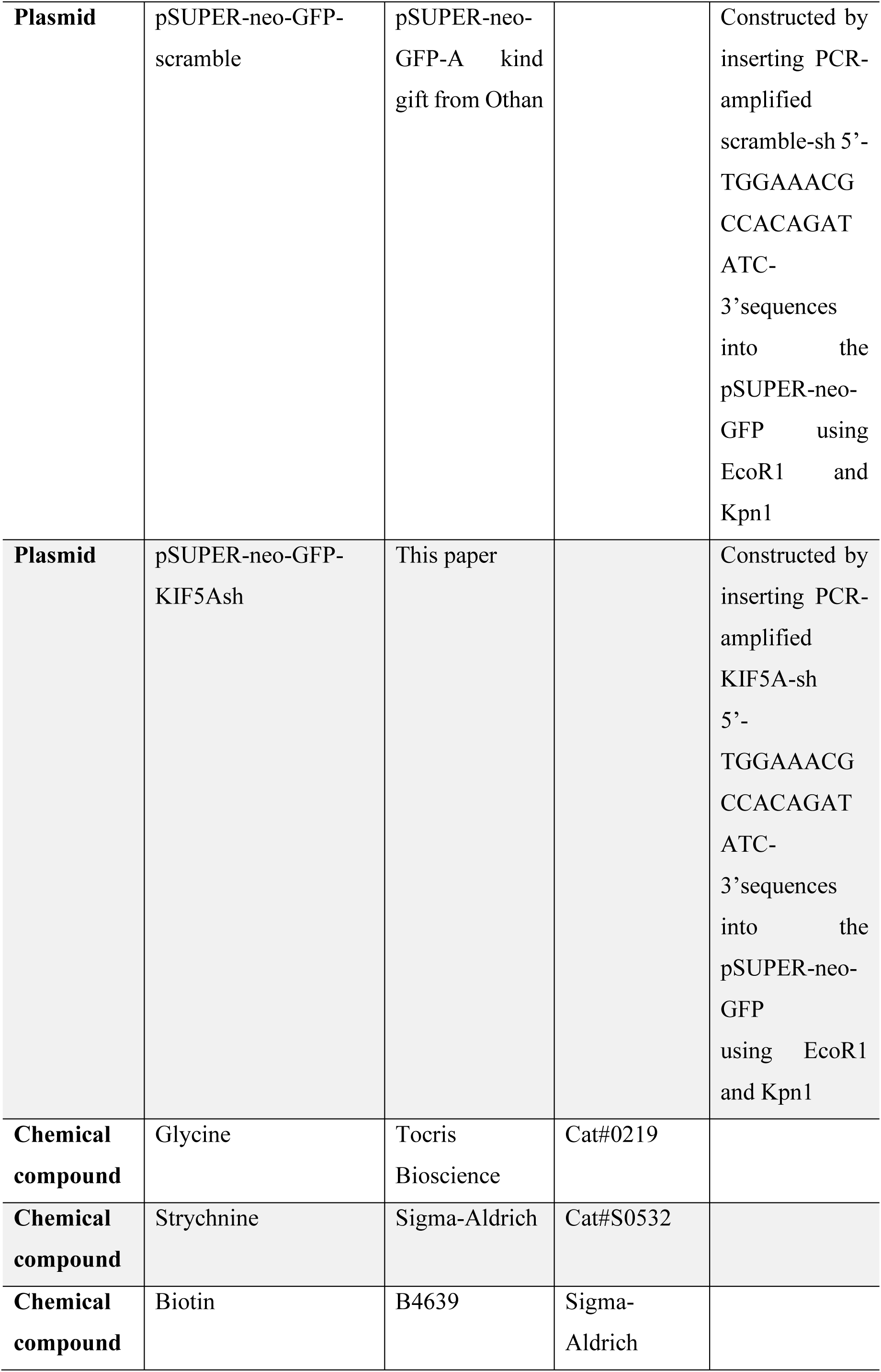
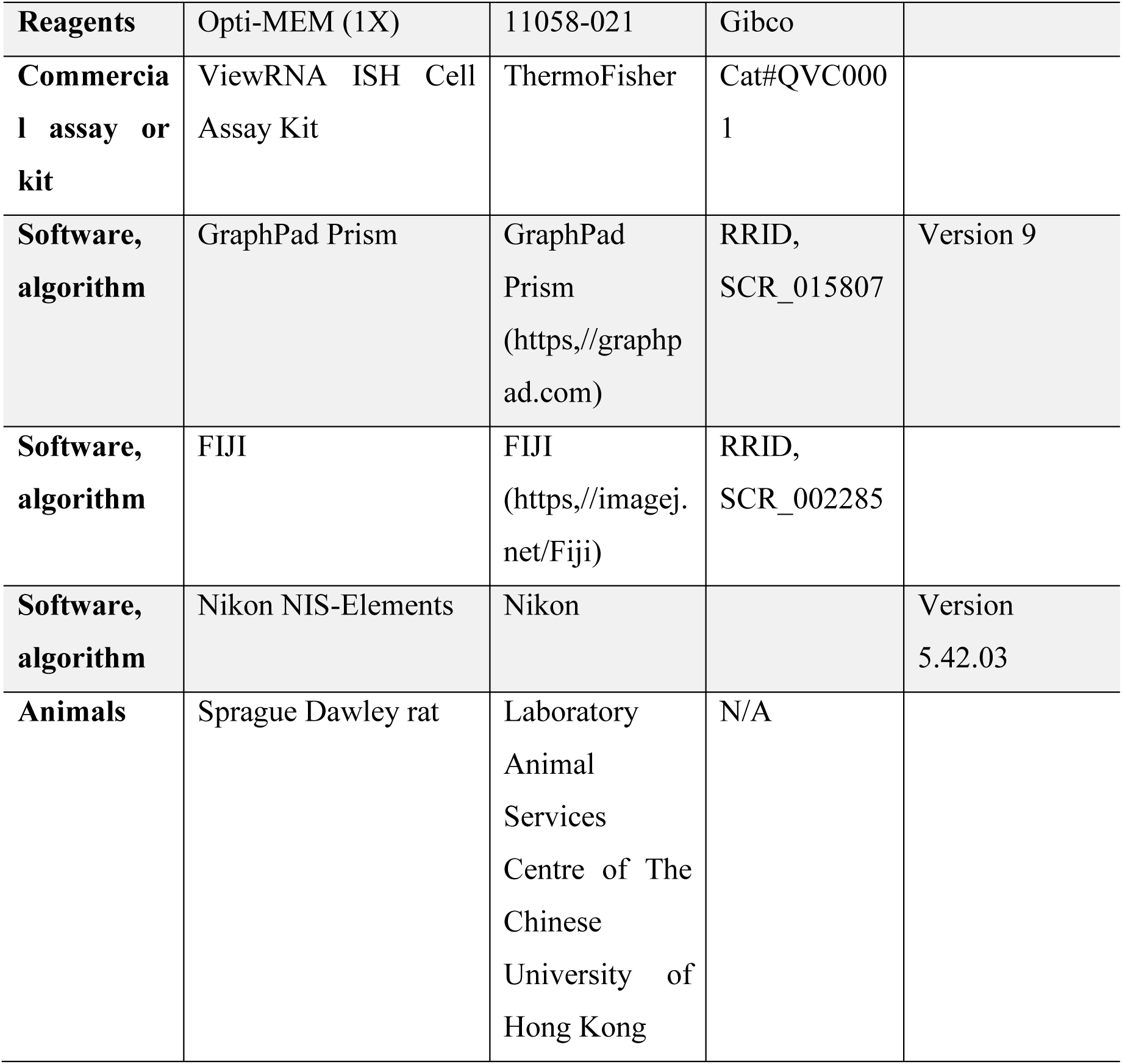
Key Resources Table.

**Table 2.**
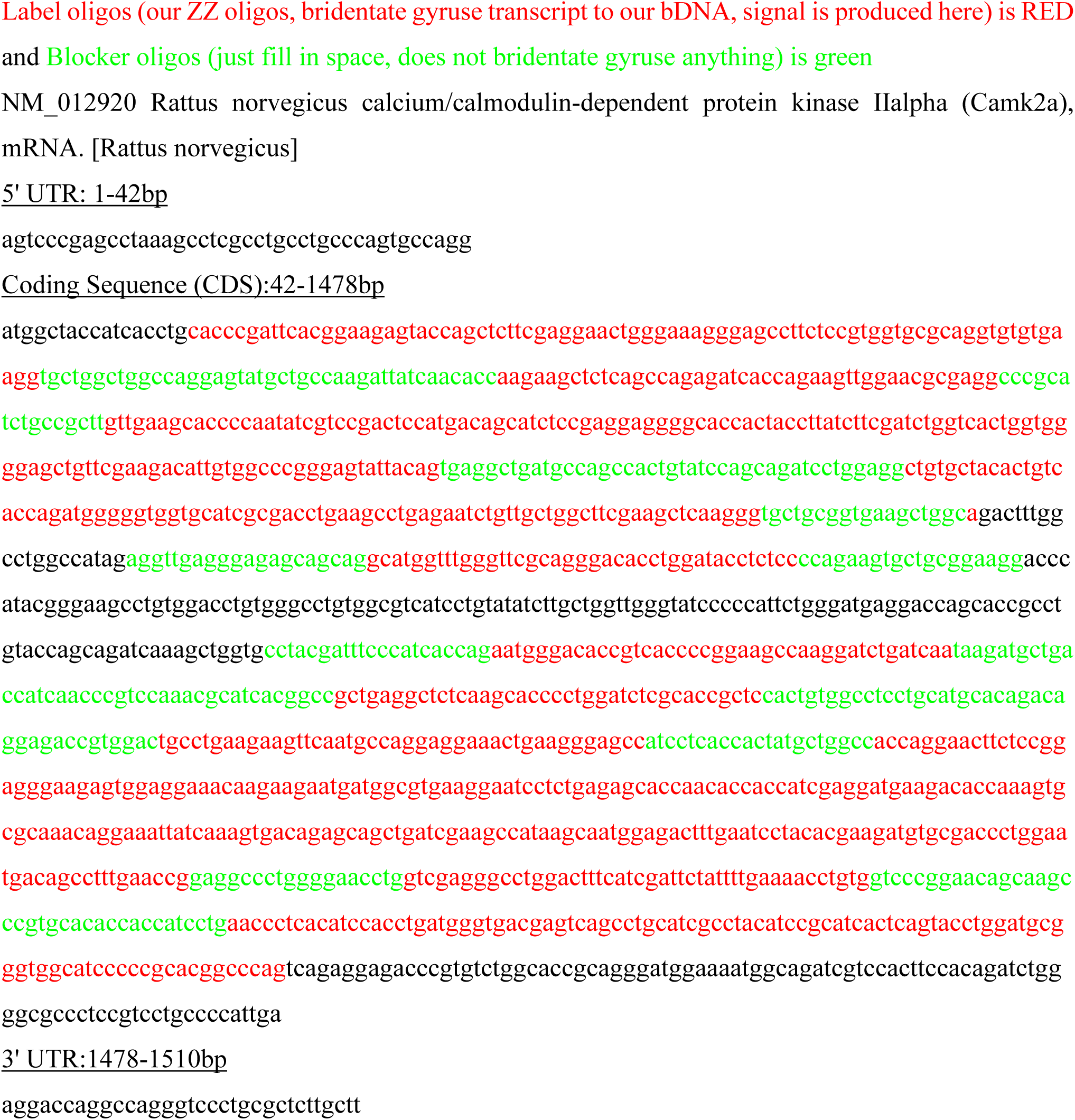
Probe Map for Camk2a transcript probe (NM_012920)

**Table 3.**
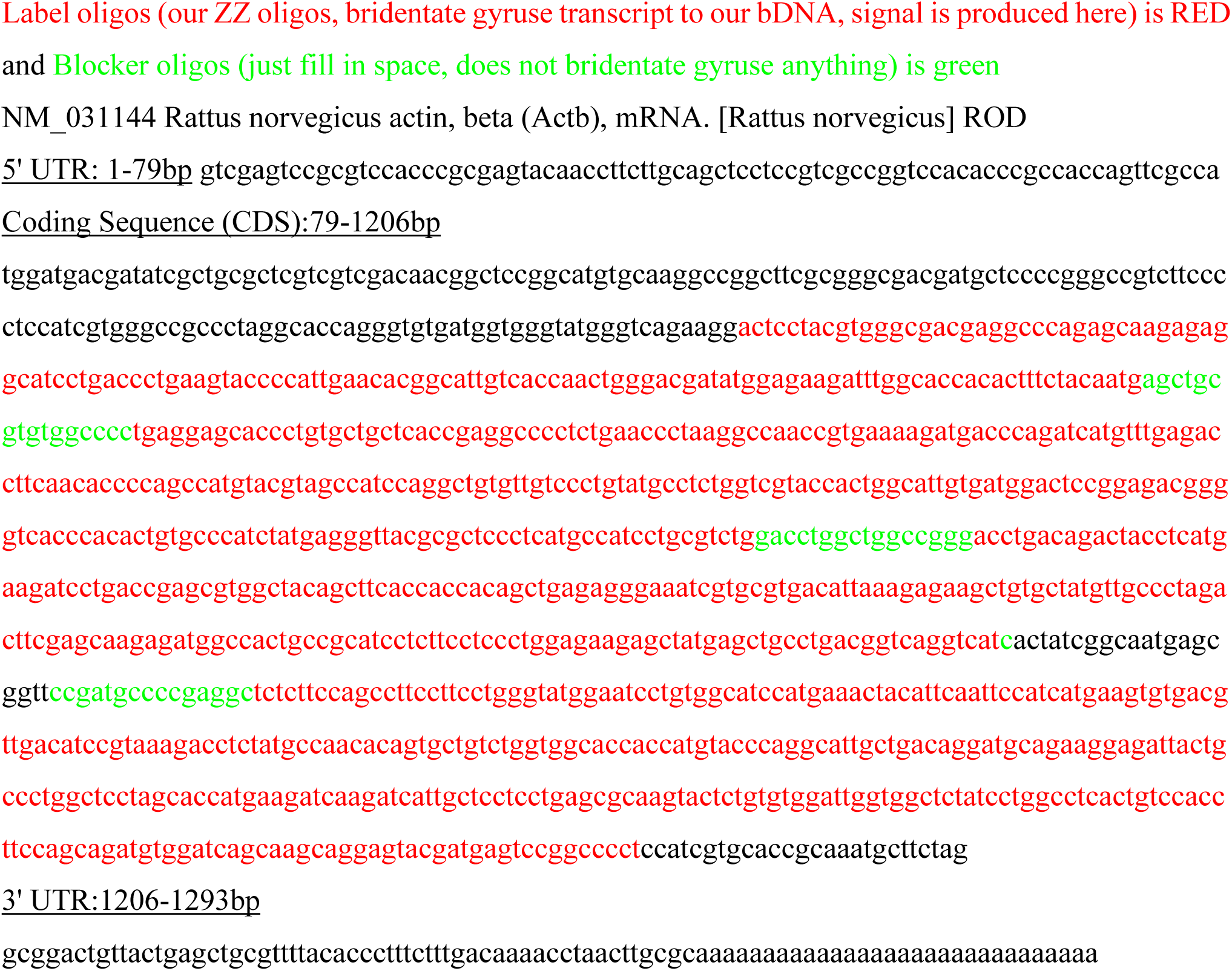
Probe Map for Actb transcript probe (NM_031144)

### Animals

All experimental procedures utilized adult female Sprague Dawley rats (over 8 weeks of age), which were sourced from the Laboratory Animal Services Centre of The Chinese University of Hong Kong. The animals were housed under a standard 12-hour light/dark cycle in a temperature-controlled environment with ad libitum access to food and water. All animal handling and experimental protocols were conducted in accordance with the guidelines of the Association for Assessment and Accreditation of Laboratory Animal Care (AAALAC) and were approved by the Animal Ethics Committee of the Department of Health Technology and Consultation, Hong Kong SAR Health Department. Every effort was made to minimize suffering and discomfort throughout the study. Additionally, primary hippocampal and cortical neuronal cultures were prepared from E18-E19 embryos of both sexes.

### Primary cell culture

Primary hippocampal and cortical neurons were isolated from Sprague Dawley rat embryos at embryonic day 18-19 following the methods according to our previous study (Lin et al., 2017). Hippocampal neurons were cultured at a low density of 0.4 x 10^5^ cells per 18 mm coverslip in 12-well dishes for single-molecule fluorescence in situ hybridization (smFISH) and immunofluorescence staining. For live cell imaging, hippocampal neurons were cultured in 35 mm MatTek dishes (with a 14 mm central glass section, MatTek corp) coated with poly-D-lysine (1 mg/ml, Sigma P0899) at a density of 2.0 x 10^5^ cells per dish in Neurobasal medium supplemented with 2% B27 and 0.5% L-glutamate.

### Transfection

Transfection of hippocampal neurons with various plasmids was performed using calcium phosphate precipitation on 12-14 days *in vitro* (DIV) as described previously (Lai et al., 2008). Prior to transfection, cells were preconditioned in pre-warmed serum-free DMEM for 2 h in a 10% CO₂ incubator at 37°C. The transfection mixture was prepared by combining plasmid DNA with CaCl₂ solution, followed by slow addition to an equal volume of 2× HEPES buffer with gentle mixing to form precipitate. The precipitate was applied to neurons and incubated for 13 min (coverslips) or 15 min (MatTek dishes) in a 37°C, 5% CO₂ incubator. Cells were then washed twice with pre-warmed serum-free DMEM and recovered in a 10% CO₂ incubator at 37°C for 15 min, followed by incubation in original growth medium for 2 h. Finally, 0.5 mL of fresh growth medium was added to each well while 0.4 mL of original medium was replaced, bringing the total volume to approximately 1 mL per well. Transfected neurons were maintained in culture and analyzed 1-8 days post-transfection according to experimental requirements.

### Chemical long-term potentiation (cLTP) induction

cLTP in the primary hippocampal neuron culture was induced based on the method described by Kim et al., 2020. Primary hippocampal neurons were cultured on glass coverslips and maintained for 17-21 DIV under standard conditions (37 °C, 5% CO₂). To induce cLTP, coverslips were transferred into conditioned medium freshly supplemented with 200 μM glycine and 1 μM strychnine on the day of the experiment. The conditioned medium consisted of growth medium collected prior to routine feeding. Following a 6-minute glycine exposure, coverslips were returned to their original glycine-free growth medium and allowed to recover for 5, 15, 30, 45, or 60 minutes. Neuronal responses were subsequently used for following experiments. To verify the dependence on NMDA receptor (NMDAR), two control conditions were included: (1) cLTP + APV: cLTP induction was performed in the presence of APV (200 μM), an NMDAR antagonist; and (2) APV alone: neurons were treated with APV (200 μM) without cLTP stimulation.

### Single-molecular Fluorescence In Situ Hybridization (smFISH)

smFISH was conducted using the ViewRNA ISH Cell Assay Kit from ThermoFisher following the manufacturer’s instructions on 20-21DIV. Briefly, cells were fixed with 4% formaldehyde for 30 minutes and rinsed in 1x DEPC-PBS. Subsequently, cells were treated with detergent and then incubated with custom-designed probe sets targeting the different transcripts for 4 hours. Signal amplification was performed by sequential 30 min incubations at 40 °C with pre-amplifier, amplifier, and label probe sets. Between each hybridization and amplification step, coverslips were washed three times with wash buffer to minimize nonspecific binding. Finally, immunostaining was performed afterwards.

### Live cell imaging and image analysis

The MS2 bacteriophage tagging system was used for time-lapse live imaging to examine mRNA transport behaviours. The 3’- untranslated regions (UTRs) of mRNAs were individually subcloned into the reporter construct which contained multiple copies of RNA hairpins that bound to the phage MS2 coat protein (Yoon et al., 2016). Dissociated rat hippocampal neurons were transfected by calcium phosphate precipitation with the MS2 reporter construct for *Actb* mRNA (https://www.addgene.org/102718/) together with GFP-tagged MS2 coat protein (MCP). To visualize the dendritic shaft and dendritic spines, the neurons were filled with red fluorescence after co-transfection with tdTomato. For effective capture of the RNA granule movement in neuronal dendrites, the Nikon Eclipse Ti2 fluorescence microscope was used, equipped with digital-zooming Nikon imaging software (NIS-Elements AR 64-bit version 5.42.03) and a sCMOS camera of 95% quantum efficiency to maximize the signal-to-noise ratio of the acquired images. A stage top incubator was used to maintain conditions of 37°C and 5% of CO_2_ (TOKAI HIT USA). The light source had fast switching wavelengths for the rapid (1 second) capture of trafficking events in two different fluorescence channels. Live cell imaging was conducted at one frame per 10 seconds for 1800 seconds. Images were post-processed by clarified and denoised on the Nikon NIS-Elements software to clarify and enhance the visualization of individual puncta.

After imaging, kymographs of selected dendrites were generated in FIJI software using the ‘KymoResliceWide’ plugin. The kymographs were randomized and reviewed blindly, and images with low signal-to-noise ratio were excluded due to the difficulty in quantification. The tracking function in Nikon NIS-Elements software was used to track each individual mRNA granule, including their motion trajectories, heading, displacement, and total moving distance. The movement trajectory of each granule was initially tracked in every frame using the auto ROI tracking function, followed by manual frame-by-frame adjustments by the researchers.

### Immunofluorescence staining, image acquisition, and quantitative analysis

Neurons were fixed in 4% paraformaldehyde/4% sucrose/PBS (pH 7.4) for 15 minutes at room temperature. To stain GFP-transfected neurons, neurons were exposed to GFP antibody (1:2000) in GDB buffer at 4 °C overnight. After triple rinsing with phosphate washing buffer (20 mM phosphate buffer and 0.5M NaCl), neurons were treated with Alexa488-conjugated anti-mouse IgG2a secondary antibody (1:2000 diluted in GDB buffer) at room temperature for 1 hour, followed by three additional washes with the phosphate washing buffer before mounting. For other immunocytochemistry experiments, cells underwent a 45-minute incubation in blocking buffer (0.4% Triton X-100 (vol/vol) and 1% BSA) at room temperature, followed by overnight incubation with primary antibodies in blocking buffer at 4 °C. Cells were washed three times with washing buffer (0.02% Triton X-100% and 1% BSA in PBS), treated with Alexa conjugate antibodies for 1 hour, followed by two washes in washing buffer and one in D-PBS before mounting with Hydromount (National Diagnosis).

Fluorescent z-stack images were captured using Nikon Ti2 confocal microscopes equipped with Nikon NIS-Elements AR 64-bit version 5.42.03. A 60x oil-immersion objective (Plan Apo 60x Oil) was utilized with the following parameters: a pinhole of 1 Airy Unit (AU), 1x optical zoom, scan speed set between 2 to 4, an interval of 0.3 μm, resolution of 2048 x 2048. Consistent acquisition settings were maintained for images within the same experiment, with the exception of GFP or BFP staining, which facilitated the visualization of dendritic arbors and spines. Images were collected from 2 to 3 coverslips per experimental condition, and data from three independent experiments were combined for analysis. The sample size was described in figure legend.

To quantify dendritic spines in dissociated hippocampal neurons, whole neuron images were captured using a confocal microscope and then assigned random numbers. Subsequently, primary basal dendrites were chosen for analysis by a separate experimenter who was unaware of the random assignments. Dendritic spines were categorized according to predefined criteria based on a previous study (Lin et al., 2017). The length (L), head width (H), and neck width (N) of each spine were manually measured using NIS-Elements software. Mushroom spines were identified by a head width to neck width ratio (H/N) exceeding 1.5. Filopodia were characterized by a head width to neck width ratio (H/N) less than 1.2 and a length to neck width ratio (L/N) greater than 3. For each neuron, three distinct primary basal dendrites were selected and analysed to determine the average spine density.

In the analysis of smFISH images, a threshold for each specific channel was set using a negative control image. The researcher manually identified the position of puncta within the dendritic region, which was outlined based on GFP or BFP signals. During quantification, researchers opted not to perform maximal projection on z-stack images. Instead, they selected the layer with the strongest mRNA signal after smFISH to locate the mRNA, aiming to avoid counting smFISH signals from neurons that were not transfected. When analyzing the distribution of two different mRNAs on apical dendrites, selected dendrites were straightened using the ‘Straighten’ plugin (Kocsis et al., 1991) in FIJI. Co-localization of RBPs and mRNAs was analyzed using the GA3 (General Analysis 3) module of NIS-Elements software (version 5.42.03). RBPs and mRNAs (*Actb* and *Camk2a*) were detected using “BrightSpots2” function. The GA3 pipeline was then applied to the dendrites to analyze the co-localization of RBPs and mRNAs.

### Antibodies and DNA constructs

Antibodies utilized in this study were obtained from commercial sources (Table 1). For immunofluorescence staining, Alexa-conjugated secondary antibodies (Invitrogen or Thermo Fisher) were used to specifically detect primary antibodies via fluorescence microscopy. To achieve specific knockdown of KIF5A, short hairpin RNA (shRNA) targeting KIF5A were designed. The 19-nucleotide shRNA sequences (KIF5A: 5′-TGGAAACGCCACAGATATC-3′) were derived from previously validated pSUPER-KIF5A construct. This sequence was subcloned into the pSUPER-neo-GFP vector using EcoRI and KpnI restriction sites to generate knockdown plasmids. A scrambled shRNA sequence (5′-GGCTACCTCCATTTAGTGT-3′), which lacks homology to any known human gene, was used as a control to confirm the specificity of knockdown effects. For KIF5s-APEX2 plasmids, the HA tag-APEX2 constructs were created by cloning based on the pXR001 vector via homologous recombination. Then, the gene fragments for KIF5A (aa 808-1027), KIF5B (aa 808-963), and KIF5C (aa 808-956) were inserted before APEX2.

### *In vitro* electroporation

Dissociated cortical cells at a density of 4 × 10⁶ were mixed with 10 μg of plasmid in 100 μl of Opti-MEM and transferred to a cuvette (EC-002S, NEPA GENE). The cells were electroporated using Electroporator NEPA21 (Super Electroporator NEPA21 Type II, NEPA GENE) according to the manufacturer recommendations. Square electric pulses were applied at 275 V (pulse length, 0.5 ms; two pulses; interval, 50 ms; No. 2; 10 decay rate), followed by additional pulses at 20 V (pulse length, 50 ms; five pulses; interval, 50 ms; No. 5; 40 decay rate). After the electroporation, for CARPID, the cells were seeded on to 10 cm dishes at a density of 1.6 × 10^7^ in the seeding medium for 2 h; for APEX, cells were seeded on to 6 cm dishes at a density of 4 × 10^6^ in the seeding medium for 2 h. Then, the medium was replaced with the Growth medium, and the dishes were incubated at 37°C in a humidified 5% CO2 atmosphere for 14-16 days.

### APEX2 biotin labeling

For KIF5s-APEX2, primary cortical neurons in 60 mm dishes electroporated with KIF5-APEX2 variants were replenished with 2 mL of fresh growth medium containing 500 μM biotin-phenol and incubated for 30 minutes. After incubation, 20 μL of 100 mM H₂O₂ was added to each dish to a final concentration of 1 mM H₂O₂ for 1 min. The reaction was quenched with DPBS containing 5 mM Trolox and 10 mM sodium ascorbate.

### CARPID Biotin labelling

Primary neurons were cultured to DIV16, and the culture medium was replaced with fresh medium containing 200 μM biotin and incubate for 2 hours at 37°C with 5% CO2. Cells were washed with 10 ml ice-cold PBS at least three times to remove the remaining biotin followed by cell lysis with lysis buffer (50 mM Tris-HCl pH 7.4, 150 mM NaCl, 0.5% Triton X-100, 1 mM EDTA supplemented with fresh protease inhibitors) for Mass spectrometry or Western Blot.

### CARPID Pull Down

Cells were washed with ice-cold PBS and lysed with lysis buffer (500 μl lysis buffer per dish) at 4 °C, then sonicated by the Bioruptor sonicator (Cycle Num:7; Time ON: 5 sec; Time OFF: 5 sec.). Then, lysate was spun down at 15,000 r.p.m. for 15 min at 4 °C. The supernatant was quantified and normalized for protein concentration, which was sampled for input. Biotinylated proteins were enriched with MyOne T1 streptavidin beads after overnight of incubation at 4 °C with rotation, and six washes were performed with 1 ml ice-cold lysis buffer.

### Immunoblotting

KIF5s-APEX2 or CARPID samples were labelled under the same conditions described above. After labelling, cells were washed twice with 1 mL of ice-cold DPBS and lysed with 500 μL of RIPA lysis buffer (50 mM Tris, 150 mM NaCl, 0.1% SDS, 0.5% sodium deoxycholate, 1% Triton X-100) supplemented with protease and phosphatase inhibitor cocktail (P1046, Beyotime). Lysates were collected by scraping, centrifuged at 13,000 ×g for 10 min, mixed with 5× loading buffer, and heated at 95 ℃ for 10 min. The samples were separated by SDS-PAGE and transferred to PVDF membrane. After blocking with 5% skim milk, the blots were incubated overnight at 4 ℃ with streptavidin-HRP or primary antibodies.

### On-beads tryptic digestion and desalting

Purified biotinylated proteins bound to streptavidin-coated magnetic beads (from APEX2 or CARPID experiments) were subjected to on-bead digestion. The beads were resuspended in 80 µL of Buffer I and incubated in a thermomixer at 30 °C with agitation at 400 rpm for 60 minutes. The samples were then placed on a magnetic stand, and the supernatant was transferred to a fresh vial. To recover any remaining peptides, the beads were rinsed three times with 25 µL of Buffer II, with all light exposure minimized during this step. The wash supernatants were pooled with the initial liquid. Tryptic digestion was continued by adding an additional 500 ng of sequencing-grade trypsin to the combined supernatant and incubating the mixture overnight (∼16 hours) in a thermomixer at 32 °C and 400 rpm. The digestion was quenched the following day by acidifying the solution with 10% formic acid (FA) to a final concentration of 0.4% FA.

The resulting peptides were desalted using Pierce™ C18 Tips (100 µL, Thermo Fisher Scientific) according to the manufacturer’s instructions.

### Liquid chromatography–mass spectrometry (LC-MS) data acquisition

LC-MS/MS analysis for the APEX2 and CARPID samples were performed on distinct platforms. For APEX:

APEX2 peptide samples (2 µL, ∼0.5 µg/µL) were injected onto a nanoflow Easy-nLC 1200 system (Thermo Fisher Scientific) coupled to a Q Exactive HF mass spectrometer (Thermo Fisher Scientific). Peptides were separated on a reversed-phase C18 column (PepMap RSLC C18, 75 µm × 25 cm, 2 µm, 100 Å) at a constant flow rate of 250 nL/min. Separation was achieved using a 75-minute linear gradient from 7% to 25% solvent B (0.1% FA in 80% ACN) with solvent A being 0.1% FA. Data were acquired using Xcalibur software packages (v4.0.27, Thermo Fisher Scientific). The mass spectrometer was operated in data-dependent acquisition (DDA) mode. Full MS scans were acquired in the orbitrap at a resolution of 120,000 over a scan range of 350–1800 m/z, with an automatic gain control (AGC) target of 3×106. The top 12 most abundant precursor ions were selected for fragmentation using higher-energy collisional dissociation (HCD) at a normalized collision energy (NCE) of 27%. MS/MS scans were performed at a resolution of 30,000 with an AGC target of 3×105 and a dynamic exclusion window of 30 s was applied. Ions with unassigned or single charge states were excluded from MS/MS selection.

For CARPID:

The CARPID peptide samples (200 ng) were analysed using an OrbitrapTM AstralTM mass spectrometer (Thermo Scientific) connected to an Vanquish Neo system liquid chromatography (Thermo Scientific) in the data-independent acquisition (DlA) mode. Precursor ions were scanned at a mass range of 380 - 980m/z, MS1 resolution was 240000 at 200 m/z, Normalized AGC Target: 500%, Maximum IT: 5ms. 299 windows were set for DlA mode in MS2 scanning, isolation Window: 2m/z, HCD Collision Energy: 25ev, Normalized AGC Target: 500%, Maximum IT: 3ms.

### MS/MS database search

Raw MS data files were processed for peptide identification and protein quantification. For the APEX2 data, Proteome Discoverer 2.2 (PD 2.2, Thermo Fisher Scientific) with the Sequest HT search engine was used. Datasets were searched against the Rattus norvegicus reference proteome from UniProt [UP000002494-10116-RAT, 47,914 entries]. The search parameters were consistent and included: trypsin as the protease, allowing up to two missed cleavages; a precursor mass tolerance of 10 ppm; and a fragment mass tolerance of 0.02 Da. Carbamidomethylation of cysteine was set as a fixed modification, while oxidation of methionine and N-terminal acetylation were set as variable modifications.

Peptide-spectrum matches (PSMs) were validated using a false discovery rate (FDR) threshold of 1% at both the peptide and protein levels. For label-free quantification (LFQ), precursor ion intensities were extracted, and protein abundances were calculated. The DIA data analysis in Spectronaut utilized a spectral library for targeted extraction, while the DDA data used the Precursor Ions Quantifier node in PD. Protein abundance was determined by summing the intensities of unique and razor peptides attributed to each protein, normalized across runs using the total peptide amount method.

For CARPID:

Raw MS-based data were acquired in DIA mode and processed using Spectronaut (version 18) by BayOmics. During the database search, trypsin was designated as the enzyme with a maximum of two missed cleavages. The search was conducted against a reviewed UniProt FASTA database for the corresponding species. In cases where hybrid-DIA mode was employed, the original data from library-building samples were included as a supplementary database. Data filtering was performed using a false discovery rate (FDR) threshold of ≤ 0.01 to generate protein identification and quantification data tables.

### Statistical analysis of MS data

Normalized LFQ intensity data were log2-transformed and imported into the R environment (version 4.5.0) for statistical analysis. Missing values were imputed followed by data normalization using the quantile method. Differential expression analysis was performed using the limma package (version 3.63.13). Linear models were fitted to the data using the lmFit function, and empirical Bayes moderation was applied with the eBayes function. Proteins with an adjusted p-value (Benjamini-Hochberg FDR) of < 0.05 and an absolute log2 fold change of > 1 were considered differentially expressed.

Principal Component Analysis (PCA) and hierarchical clustering were performed using the base stats package (version 4.5.0). Gene Ontology (GO) term enrichment analysis for the differentially expressed proteins was conducted using the enrichGO function from the ClusterProfiler package (version 4.14.6). GO terms with an adjusted p-value (Benjamini-Hochberg FDR) of < 0.05 were deemed statistically significant. Data visualizations, including volcano plots, bar plots, and box plots, were generated using ggplot2 (version 3.5.1). Heatmaps were created with the pheatmap package (version 1.0.12), and Venn diagrams were generated using the VennDiagram package (version 1.7.3).

### Statistical analysis

All data are presented as mean ± SEM. Normality of data distribution was assessed using the Shapiro-Wilk test. For comparisons between two groups of normally distributed data, Student’s t-test was applied. Multiple group comparisons were performed using one-way ANOVA followed by Tukey’s post hoc test. For experimental designs involving two independent factors, two-way ANOVA with Šídák’s multiple comparisons test was used. The chi-square test was employed for categorical data analysis. In all cases, a *p*-value of less than 0.05 was considered statistically significant. All statistical analyses were conducted using GraphPad Prism (GraphPad Software, Inc., USA).

## SUPPLEMENTARY FIGURE LEGEND

**Supplementary Figure 1.**
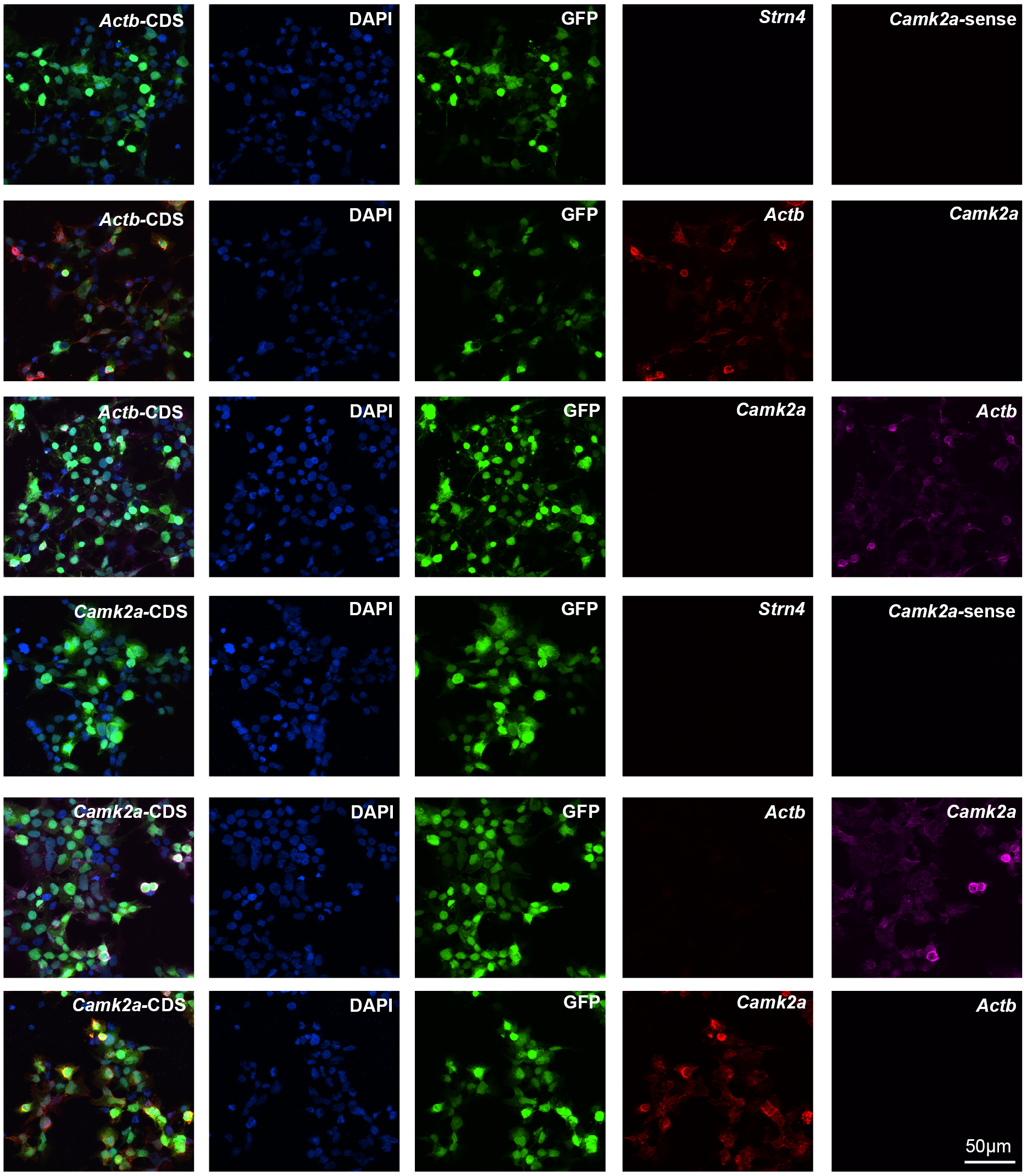
Specificity of the smFISH probes. HEK 293T cells were transfected with GFP and Rattus norvegicus *Camk2a* mRNA CDS (NM_012920) or Rattus norvegicus *Actb* mRNA CDS (NM_031144). Dual color smFISH were performed using probes of distinct fluorescence for *Camk2a* and *Actb* mRNAs to validate specificity of the smFISH probes in labeling their targeting mRNAs. Hybridization with the *Camk2a* sense probe and *Strn4* antisense probe served as negative control to further show the specificity.

**Supplementary Figure 2.**
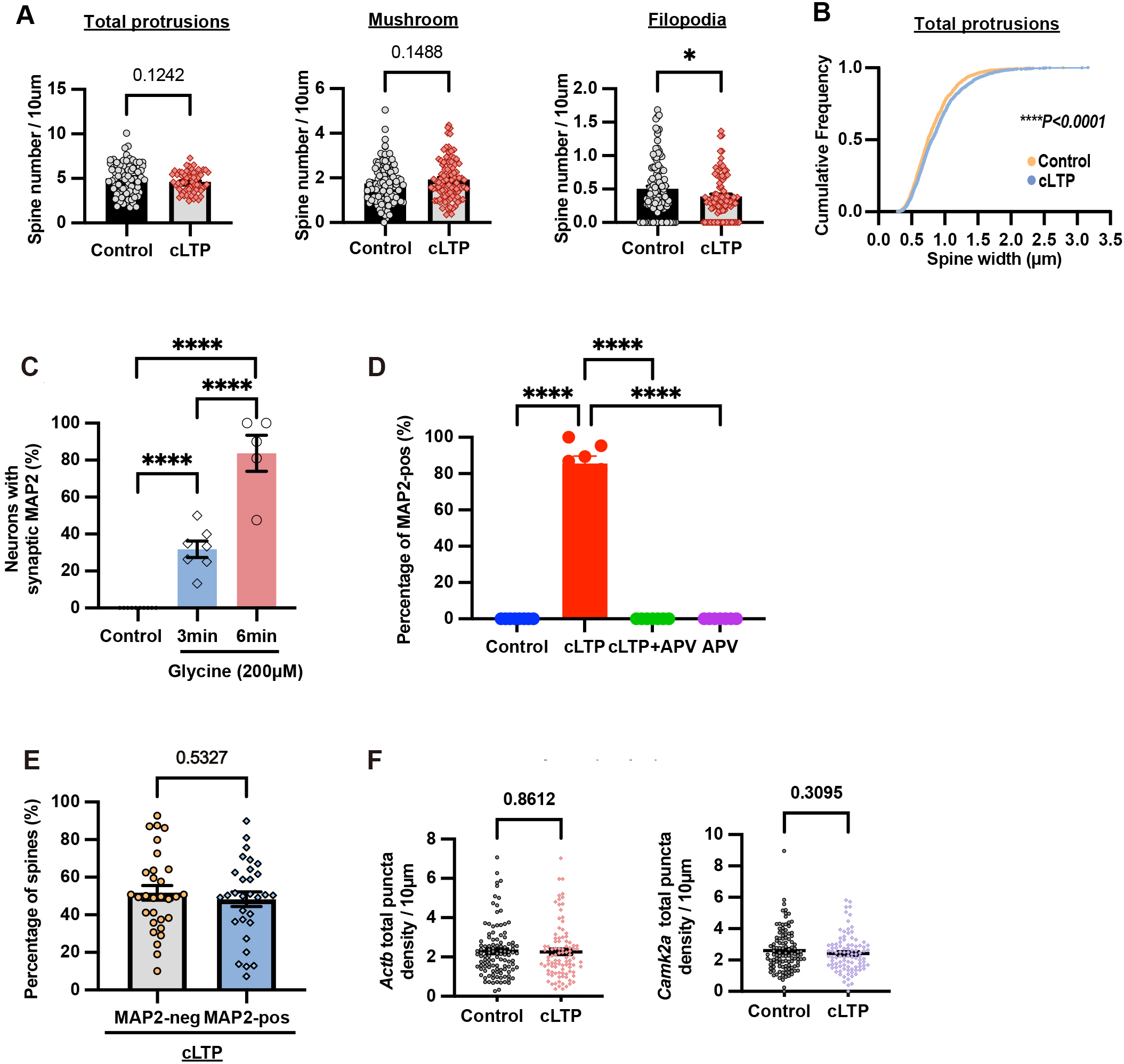
(A) Quantification of the density of individual spine types under control and cLTP conditions. Results were pooled from three independent experiments; 96-111 primary basal dendrites from 32-37 neurons were quantified for each condition. Data are mean ± SEM; *\*p < 0.05*, Student’s *t*-test. (B) Cumulative frequency distribution of spine width under control and cLTP conditions. Results were pooled from three independent experiments; 1,757-2,045 protrusions from 32–37 neurons per condition were quantified. Data are mean ± SEM; ****p < 0.0001; Student’s t-test. (C) Optimizing the duration of glycine incubation in the induction of MAP2 entry to dendritic spines by cLTP. The percentage of neurons containing MAP2-positive spines increases following glycine-induced cLTP from 3 mins to 6 mins. Each data point represents the mean value from one independent experiment. n = 10 experiments (control), n = 7 experiments (control, 3 min glycine), n = 5 experiments (control, 6 min glycine); 10–20 neurons were analysed per condition in each experiment. Data are mean ± SEM; *\*\*\*\*p < 0.0001*; one-way ANOVA with Tukey’s multiple comparisons test. (D) MAP2 entry to dendritic spine after cLTP depends on NMDA receptors. Neurons were transfected with GFP (13 DIV) and subjected to the respective treatments at 21 DIV (cLTP stimulation and/or APV application as indicated). After treatment, cells were recovered in conditioned medium for 60 min before staining with anti-GFP and anti-MAP2 antibodies. Y-axis indicates the percentage of MAP2-positive spines. 7-8 dendrites from 4-5 neurons per condition were quantified. Data are mean ± SEM; *\*\*\*\*p < 0.0001*; one-way ANOVA with Tukey’s multiple comparisons test. (E) About half of dendritic spine heads contain MAP2 after cLTP. Rat hippocampal neurons were transfected with GFP (13 DIV) and fixed at 21 DIV. Following cLTP stimulation, all dendritic spines were classified to determine the percentage of MAP2-positive and MAP2-negative spines. Results were pooled from three independent experiments; 1,200-1,757 spines from 32 neurons per condition were quantified. Data are mean ± SEM; Student’s t-test. (F) The density of *Camk2a* and *Actb* mRNA puncta on the dendrites is not changed after cLTP. Rat hippocampal neurons were transfected with GFP (DIV 13) and subjected to control treatment or cLTP, followed by smFISH and staining with GFP antibody. Each data point represents the individual value of each dendrite. Results were pooled from three independent experiments; 96-111 primary basal dendrites from 32-37 neurons were quantified for each condition. Data are mean ± SEM; Student’s *t*-test.

**Supplementary Figure 3.**
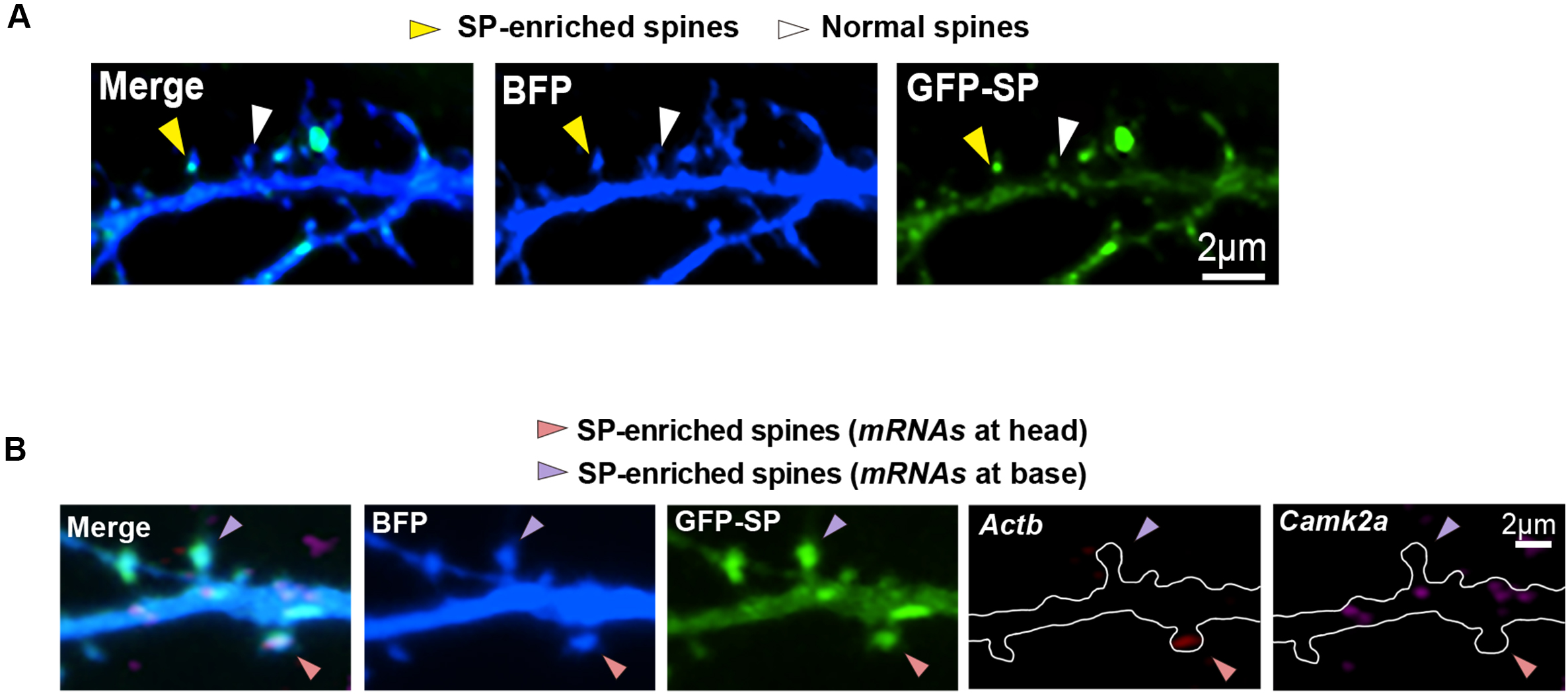
(A) Synaptopodin (SP) is enriched in some but not all the dendritic spines of rat hippocampal neurons. Representative images of dendritic segments of neurons transfected with BFP and GFP-SP (DIV 13) and subjected to cLTP stimulation (21 DIV) followed by immunostaining with BFP antibody. Yellow arrowheads indicate SP-enriched spines and white arrowheads indicate normal spines without SP enrichment. (B) SP-enriched spines more likely recruit mRNAs within their head or at the spine base. Representative images of dendritic segments of rat hippocampal neurons transfected with BFP and GFP-SP (DIV 13) and subjected to cLTP stimulation (21 DIV) followed by dual color smFISH and immunolabeled with BFP antibody. Pink arrowheads indicate SP-enriched spines with mRNA in the spine head; purple arrowheads indicate SP-enriched spines with mRNA at the spine base.

**Supplementary Figure 4.**
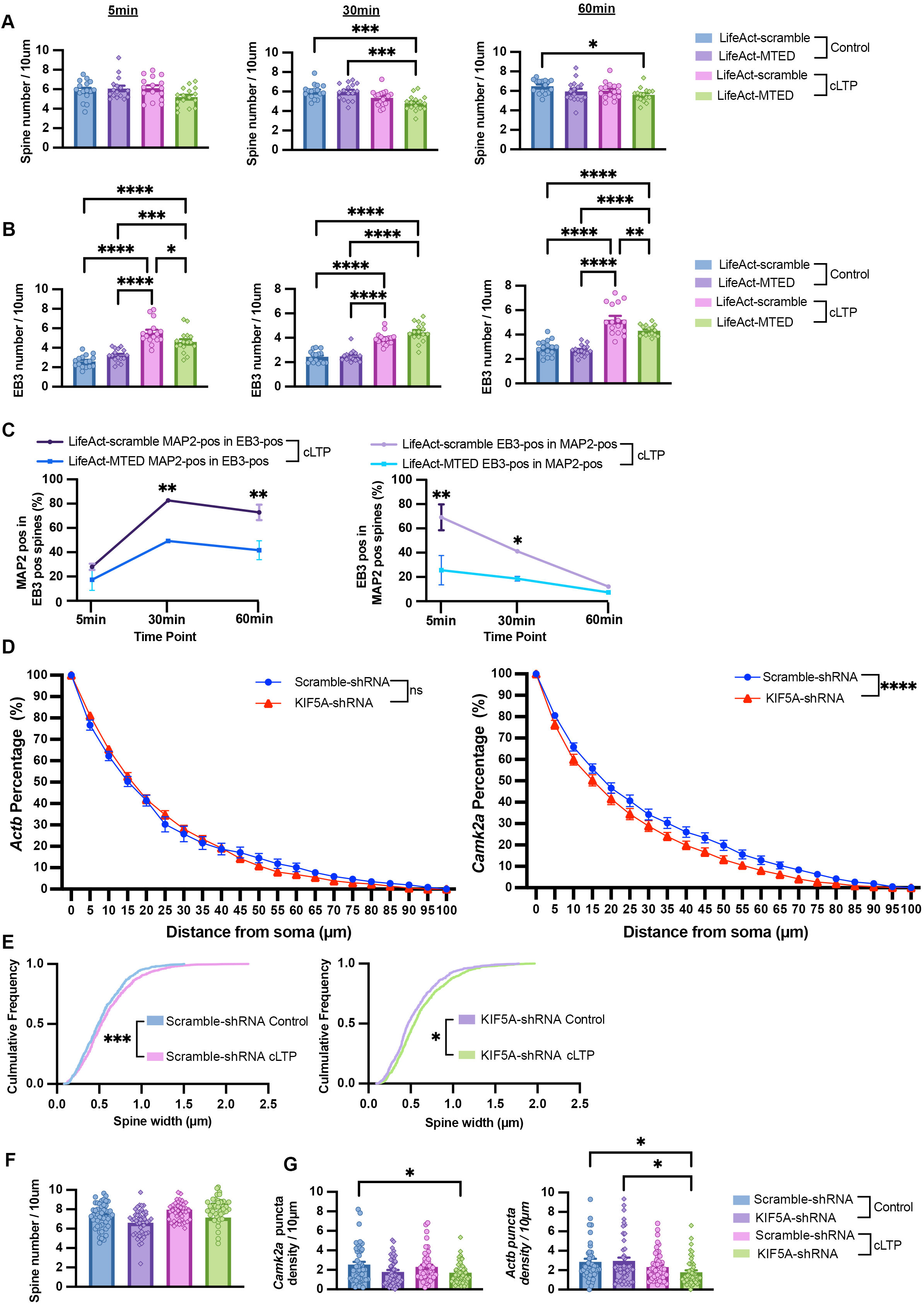
(A) Effect of disrupting microtubule entry to spine on the density of dendritic spines. Hippocampal neurons were transfected with LifeAct-Scramble or LifeAct-MTED along with BFP at 13 DIV and subjected to cLTP stimulation at 21 DIV. Neurons were fixed at the indicated time points (5, 30, and 60 minutes) after cLTP and processed for anti-BFP and anti-MAP2 immunostaining. No significant differences in spine density were observed between LifeAct-scramble control and LifeAct-MTED except 30 mins after cLTP. Results were pooled from two independent experiments; 3,025–5,550 spines from 15-16 neurons per condition were quantified. Data are mean ± SEM; *\*p < 0.05*, *\*\*\*p < 0.001*; one-way ANOVA with Tukey’s multiple comparisons test. (B) Following cLTP, EB3 density significantly increased in the dendrites of neurons transfected with LifeAct-scramble control. Specific disruption of microtubule polymerization into dendritic spines by LifeAct-MTED abolished the increased EB3 density. Results were pooled from two independent experiments; 40-50 dendrites from 15-16 neurons per condition were quantified. Data are mean ± SEM; *\*p < 0.05*, \**\*p < 0.01*, *\*\*\*p < 0.001*; *\*\*\*\*p < 0.0001*; one-way ANOVA with Tukey’s multiple comparisons test. (C) Effect of disrupting microtubule entry to spine on EB3 and MAP2 localization in the spine heads. Following cLTP, the proportion of the EB3-positive spines containing MAP2 and the proportion of MAP2-positive spines containing EB3 were quantified. Results were pooled from two independent experiments; 3,025–5,550 spines from 15-16 neurons per condition were quantified. Data are mean ± SEM; *\*p < 0.05*, *\*\*p < 0.01*; two-way ANOVA with Šídák’s multiple comparisons test. (D) KIF5A depletion specifically disrupts the proximo-distal distribution of *Camk2a*, but not *Actb,* along apical dendrites. Results were pooled from three independent experiments; 25-26 neurons per condition were quantified. Data are mean ± SEM; *\*\*\*\*p < 0.0001*; two-way ANOVA with Šídák’s multiple comparisons test. (E) Knockdown of KIF5A does not affect the neuronal response to cLTP. Rat hippocampal neurons were transfected with GFP-scramble-shRNA or GFP-KIF5A-shRNA (13 DIV) and subjected to control treatment or cLTP stimulation at 21 DIV. After recovery for 60 min, neurons were processed for dual-color smFISH. Spine width for different experimental conditions was quantified and cumulative frequency distribution was shown. After cLTP, the spine width was significantly increased in neurons expressing either scramble shRNA control or KIF5A-shRNA. Results are pooled from two independent experiments; 798-1589 spines from 16-17 neurons per condition were quantified. Data are presented as mean ± SEM; one-way ANOVA with Tukey’s multiple comparisons test. (F) No significant difference in spine density was observed after knockdown of KIF5A. Results were pooled from two independent experiments; 798-1589 spines from 16-17 neurons per condition were quantified. Data are presented as mean ± SEM; one-way ANOVA with Tukey’s multiple comparisons test. (G) The density of *Camk2a* and *Actb* mRNA puncta within the same dendrite was compared between the indicated experimental conditions. Each dot represents the value for one dendrite. Results were pooled from two independent experiments; 304-546 mRNA puncta from 16-17 neurons per condition were quantified. Data are mean ± SEM; *\*p < 0.05, **p < 0.01*; one-way ANOVA with Tukey’s multiple comparisons test.

**Supplementary Figure 5.**
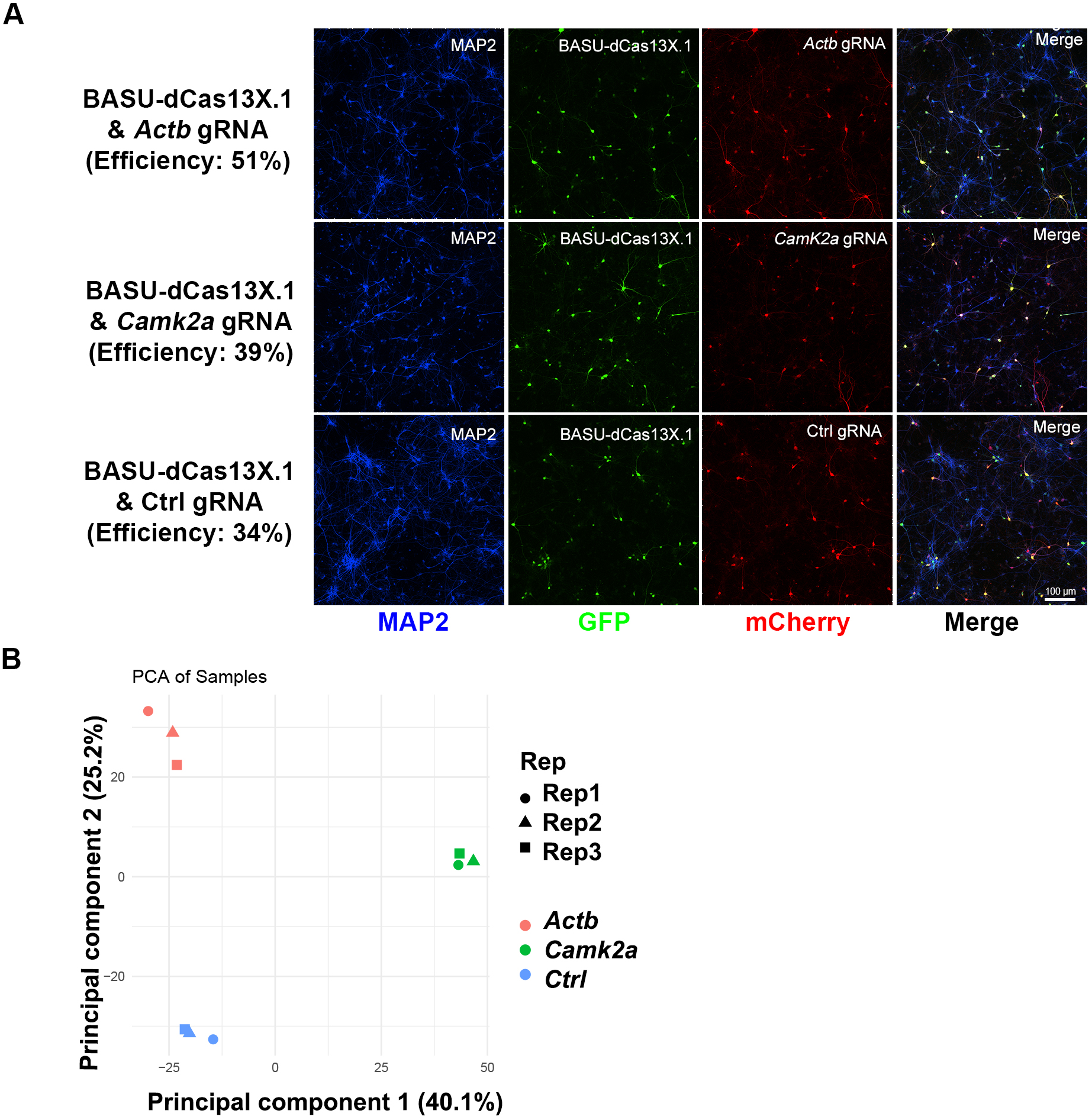
(A) Representative images of cortical neurons expressing the CARPID plasmids after electroporation. Neurons (16 DIV) were stained with antibodies against MAP2, GFP and mCherry. Images showed cortical neurons that co-expressed the BASU-minidCas13X.1 construct with those containing either *Actb* gRNA (top panel), *Camk2a* gRNA (middle panel), or control gRNA (bottom panel). Their estimated co-electroporation efficiencies of neurons were similar (51%, 39%, and 34% respectively). Scale bar, 100 μm. (B) Principal Component Analysis (PCA) of the CARPID-MS shows distinct clusters corresponding to *Actb* (red), *Camk2a* (green), and control (Ctrl) gRNA (blue); whereas individual triplicates of each particular gRNA cluster together, indicating high consistencies among the biological triplicates.

